# The adaptive immune response to *Trichuris* in wild versus laboratory mice: An established model system in context

**DOI:** 10.1101/2023.08.28.555155

**Authors:** Iris Mair, Jonathan Fenn, Andrew Wolfenden, Ann E. Lowe, Alex Bennett, Andrew Muir, Jacob Thompson, Olive Dieumerci, Larisa Logunova, Susanne Shultz, Janette E. Bradley, Kathryn J. Else

## Abstract

Laboratory model organisms have provided a window into how the immune system functions. An increasing body of evidence, however, suggests that the immune responses of naive laboratory animals may differ substantially to those of their wild counterparts. Past exposure, environmental challenges and physiological condition may all impact on immune responsiveness. Chronic infections of soil-transmitted helminths impose significant health burdens on humans, livestock and wildlife, with limited treatment success. In laboratory mice, Th1 versus Th2 immune polarisation is the major determinant of helminth infection outcome. Here we compared antigen-specific immune responses to the soil-transmitted whipworm *Trichuris muris* between controlled laboratory and wild free-ranging populations of house mice (*Mus musculus domesticus*). Wild mice harbouring chronic, low-level infections produced lower levels of cytokines in response to Trichuris antigen than laboratory-housed C57BL/6 mice. Wild mouse effector/memory CD4+ T cell phenotype reflected the antigen-specific cytokine response across the Th1/Th2 spectrum. Increasing worm burden was associated with body condition loss. However, local Trichuris-specific Th1/Th2 balance was positively associated with worm burden only in older wild mice. Thus, although the fundamental relationships between the CD4+ T helper cell response and resistance to T. muris infection are similar in both laboratory and wild *M.m.domesticus,* there are quantitative differences and age-specific effects that are analogous to human immune responses. These context-dependent immune responses demonstrate the fundamental importance of understanding the differences between model and natural systems for translating mechanistic models to ‘real world’ immune function.

## Introduction

The immune system is a highly complex and adaptable system, shaped throughout a lifetime by lifestyle and environmental factors. Unsurprisingly, the immune system of ‘naïve’ adult laboratory mice resembles that of human newborns more than that of human adults (1). It is therefore important to cross-validate laboratory models within ecologically relevant frameworks, and delineate associations in complex, natural systems that may not be possible to be recapitulated in the laboratory setting (2). Indeed, there are now multiple reports pointing to wider environmental influences being critical in determining infection outcome and health impacts of experimental infections (3, 4).

Measurement of functional adaptive immune responses in humans or wild animals is challenging. In humans, there are obvious ethical constraints limiting sampling to minimally-invasive studies, and human studies are logistically and financially particularly challenging. For wild non-model species, immunological analysis tools are often limited (5–7), and ethical considerations can limit study designs. Despite these difficulties, several studies have been performed recently to understand variation in T cell immune responses in wild mammals (8–13). Further, pioneering studies have started to probe immune responses to parasite infection in *M. m. domesticus* in the wild (14) and laboratory mice released into outdoor enclosures (4). However, no study in wild mammalian populations has quantified the local parasite-specific CD4^+^ T cell response which is deemed key to infection outcome, its association with infection in an uncontrolled setting, how the wild parasite-specific immune response compares to laboratory models currently used for biomedical research, and what the ultimate ecological consequences are for a given host animal.

We present a study that, for the first time, probes the local immune response to the Soil-Transmitted Helminth (STH) *Trichuris muris* in a naturally infected wild host and the potential health implications of individual variation in infection-immunity dynamics. STHs affect virtually all mammalian species, including humans, livestock and wildlife. Both health and economic concerns drive research into understanding disease susceptibility and potential ways to minimise disease burden both on the individual and population level. By every measurable health statistic, low/low-middle income countries are disproportionately affected by STH, which include the gut-dwelling nematode parasite *Trichuris trichiura* (15). Studies of peripheral immune responses to STH in humans reveal a complex picture, with low dose chronic infections the norm and a potential build-up of Th2-mediated immunity after decades of infection exposure (16, 17). Infections are typically overdispersed within a population (18, 19), an epidemiological pattern mirrored in wild rodents (20).

In order to better understand host-pathogen interactions and to drive novel therapeutic development, experimental work in laboratory mouse models has been employed for several decades. A particularly useful helminth model species is *T. muris*, a natural parasite of *M. musculus,* which has high genome conservation with *T. trichiura* (21, 22). Laboratory studies employ largely single dose infections, often classified as either high dose (>100 eggs) or low dose (<40 eggs), in highly controlled environments. Such laboratory-based systems have demonstrated the importance of Th2 responses for the successful expulsion of gastrointestinal (GI) nematodes (16, 23, 24). Subsequent studies in the laboratory mouse model have gone on to show that environmental context and host-associated factors are important in determining the quality of the anti-parasite immune response and thus infection outcome. For example, infection regime (25), diet (26, 27), sex (28) and age (29) each play a significant role in shaping resistance to infection in the laboratory setting.

To allow in-depth immunological analysis, coupled with natural ecological variation, we studied a fully wild and isolated island population of house mice living with minimal human interference, using longitudinal and cross-sectional sampling. Given the multitude of host-intrinsic and -extrinsic factors at play in the wild, we hypothesised that factors shown to determine the immune response and susceptibility to *Trichuris* infection in the lab would have less of an impact in the wild. However, we further hypothesised that despite the ‘noise’ of this uncontrolled study system, we would be able find evidence of a link between T helper cell polarisation and disease susceptibility. In this study, we show that the local recall cytokine response is quantitatively significantly weaker in wild mice compared to those in a laboratory experimental setting, but showed similarities in the Th1/Th2 balance of their recall cytokine response. Of several parameters measured, only age and worm burden explain, to a small degree, local anti-parasite cytokine responses. Importantly, only in older, but not younger wild mice, did we find the evidence of a link between T helper cell polarisation and worm burden. Furthermore, longitudinal data analysis revealed a connection between changes in infection levels and host health. Overall, our study population mimicked several parasite-immune relationships found in human populations where Trichuris is endemic; and highlights that the uncontrolled environment likely supersedes several of the mechanisms found in laboratory settings. We suggest that by studying the immune response in a wild house mouse system alongside more controlled and mechanistic laboratory studies, we can expand our well-established laboratory disease models to include an ecologically relevant context and thereby enhance their utility for translational biomedical research.

## Results

### The Isle of May mice harbour chronic *Trichuris muris* infections, presenting a measurable health impact at high infection levels

Worm burden was assessed upon cull of 272 wild mice, revealing a *Trichuris* prevalence of 79%. ITS1-5.8S-ITS2 rDNA sequencing confirmed the parasite species to be *T. muris*, with 100% sequence overlap with the Edinburgh (or E) isolate used in laboratory studies for decades (30) (Suppl. Figure 1), and closely associating with other Western European isolates previously reported in wild-caught rodents (31) (Suppl. Figure 2). The majority of mice harboured low levels of *Trichuris* (<10 worms) (Figure 1A), in an overdispersed distribution pattern, with mean and median worm counts of infected individuals being 11.5 and 5, respectively. Animals without worms but seropositive for anti-Trichuris IgG1/IgG2a represented 20% of the study population. Only 1% of animals were both free of worms and seronegative. Although the majority of animals that were visually classified as juveniles already presented with worms, the prevalence of worm infections in adults was significantly higher than in juveniles (Figure 1B). Using a logistic regression model and our proxy for age – maturity index (see methods for details), we confirmed that older animals had a significant higher chance of being infected than younger mice (linear regression estimate 0.369, *p*=2.13×10^−4^). Furthermore, maturity index was the only factor positively associated with worm burden in a generalised linear model (GLM) (Figure 1C, see Suppl. Table 1 for full model output), i.e. animals tend to accrue rather than expel worms with increasing age. Overall, these data support the hypothesis that exposure to *Trichuris* infection is high in this wild mouse population, and infections are chronic in nature.

**Figure 1:**
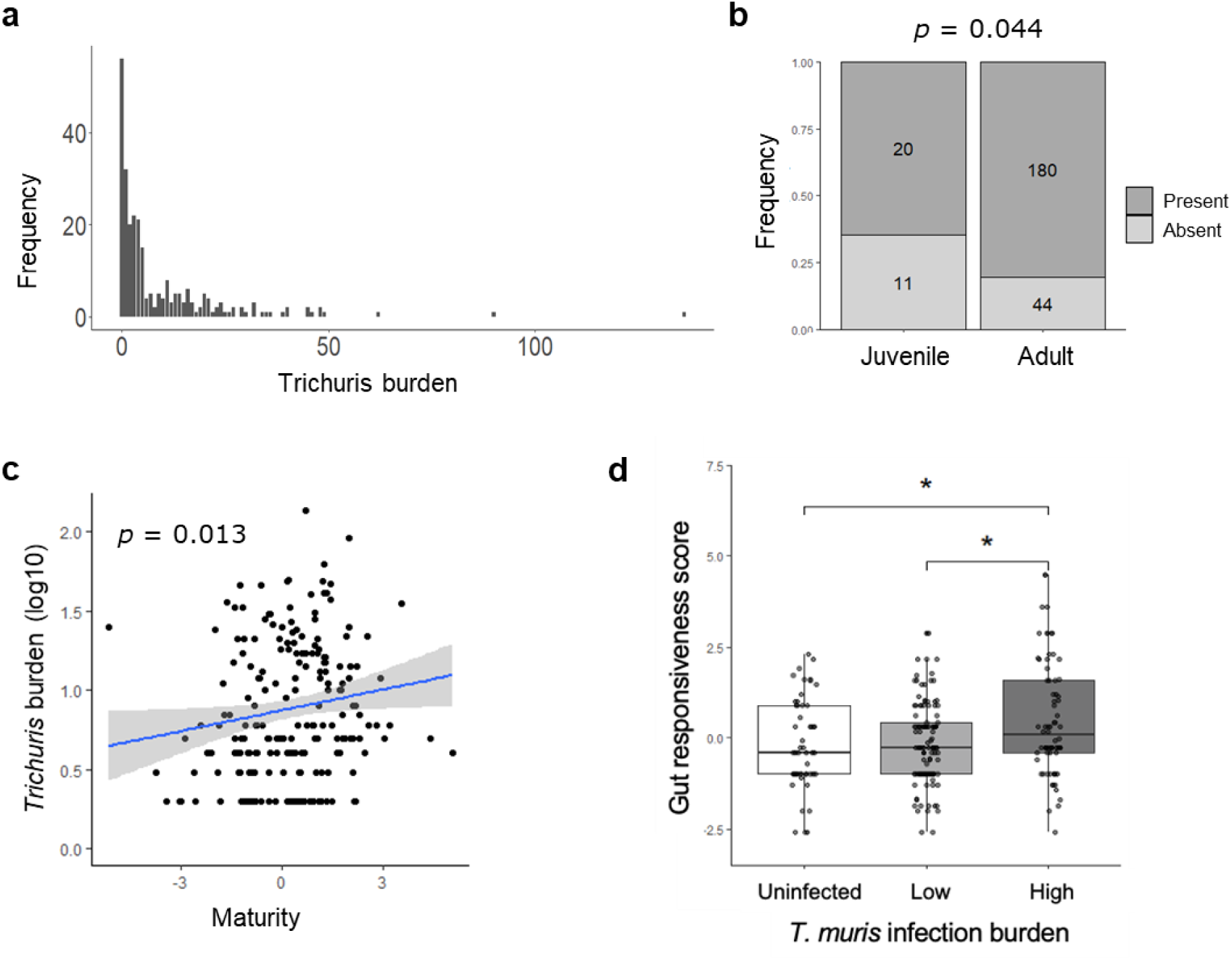
*Trichuris muris* is a highly prevalent endoparasite in Isle of May mice causing mainly chronic infection. **a)** Frequency distribution of *Trichuris* burden in wild caught *Mus musculus* on the Isle of May, assessed microscopically *post-mortem*. **b)** Prevalence of Trichuris infection in juveniles and adults, analysed using a Pearson’s Chi-squared test. **c)** Correlation plot of Trichuris burden and maturity index as proxy for age, with p value derived from a generalised linear model explaining Trichuris burden with maturity index, sex, date of capture and scaled mass index (accounting for year of capture). **d)** Group-wise comparison of gut responsiveness score between uninfected individuals, low level infected individuals (1-9 worms) and higher level infected individuals (>9 worms) (Kruskal-Wallis test followed by a Dunn’s test). *p≤0.5, **p≤0.01, ***p≤0.001.

As is the case with many free-living animals and affected human populations, the Isle of May mouse population was polyparasitised. In addition to *T. muris*, two further helminth endoparasites were found as described in a previous study (32). Specifically, the gut-dwelling pinworm *Syphacia obvelata* and the liver nematode *Calodium hepaticum* were found in a number of animals over the duration of this study with a low prevalence of *S. obvelata* and high prevalence of *C. hepaticum* (Suppl. Figure S3). To assess the health consequences of harbouring *T. muris* at the organ level, we generated a proxy for gut architecture disturbance which we termed ‘gut responsiveness’ (see methods and Suppl. Figure S4 for details). *Trichuris* burden was positively associated with an increase in gut responsiveness at worm burdens of 10 or higher (Kruskal-Wallis *p*=0.023), suggesting that infection level is an important determinant of the impact of infection on host health (Figure 1D).

### The cytokine recall response to *Trichuris* in wild mice is significantly weaker than in laboratory single dose infection models

We next quantified the local adaptive immune responses to parasite exposure of wild mice living in a natural environment versus those of laboratory mice. To this end, upon dissection, mesenteric lymph node cells of wild mice and experimentally infected laboratory mice underwent recall stimulation with parasite excretory-secretory product (E/S) and secreted cytokines were measured (Figure 2A-C). A principal component analysis (PCA) was applied to wild mouse data to reduce the complexity of the multi-cytokine response (Figure 2D). Principal component 1 (PC1), capturing 42% of the variation in cytokine expression, denoted increasing cytokine concentrations overall. Whilst no discrete clusters were apparent within the wild mouse population, the loadings of PC2 (18% of variation) clearly pointed towards animals shifting along an axis of either a more Th2-dominated (IL-4, IL-5, IL-9, and IL-13) or a regulated Th1/pro-inflammatory response (TNF-a, IL-6, IFN-g, IL-17, IL-10). Strikingly, cytokine responses did not differ between wild mice harbouring *Trichuris* worms or not, at the time of analysis, in the context of both response strength (PC1) and response quality (PC2) (Figure 2D). Inclusion of laboratory animals in the analyses revealed that the cytokine responses of the majority of wild and laboratory animals overlapped at least partially (Figure 2E), with the notable exception of laboratory mice culled at day 34 post high dose infection, at a time point when worms have likely been expelled under this experimental regime. Wild mice overall secreted significantly lower levels of cytokines upon recall than laboratory mice under any of the infection regimes and time points (Figure 2F). For reference, naive lab mice were included as controls in a separate PCA and exhibited a lower median for overall cytokine expression in response to *Trichuris* E/S (PC1 median: −1.3193, IQR: −2.4500, −0.3607) compared to wild mice (PC1 median: −0.2544, IQR: −1.3020, 0.8524). PC2, or the Th1/Th2 balance of the parasite-specific immune response, in wild animals overlapped fully with C57BL/6 mice at day 21 and day 34 following a low dose infection. In summary, whilst the Th1/Th2 balance of the anti-*Trichuris* cytokine response in wild mice was comparable to a laboratory chronic infection model, mLN cells of wild mice overall secreted less cytokine in response to recall stimulation with *Trichuris* E/S product.

**Figure 2:**
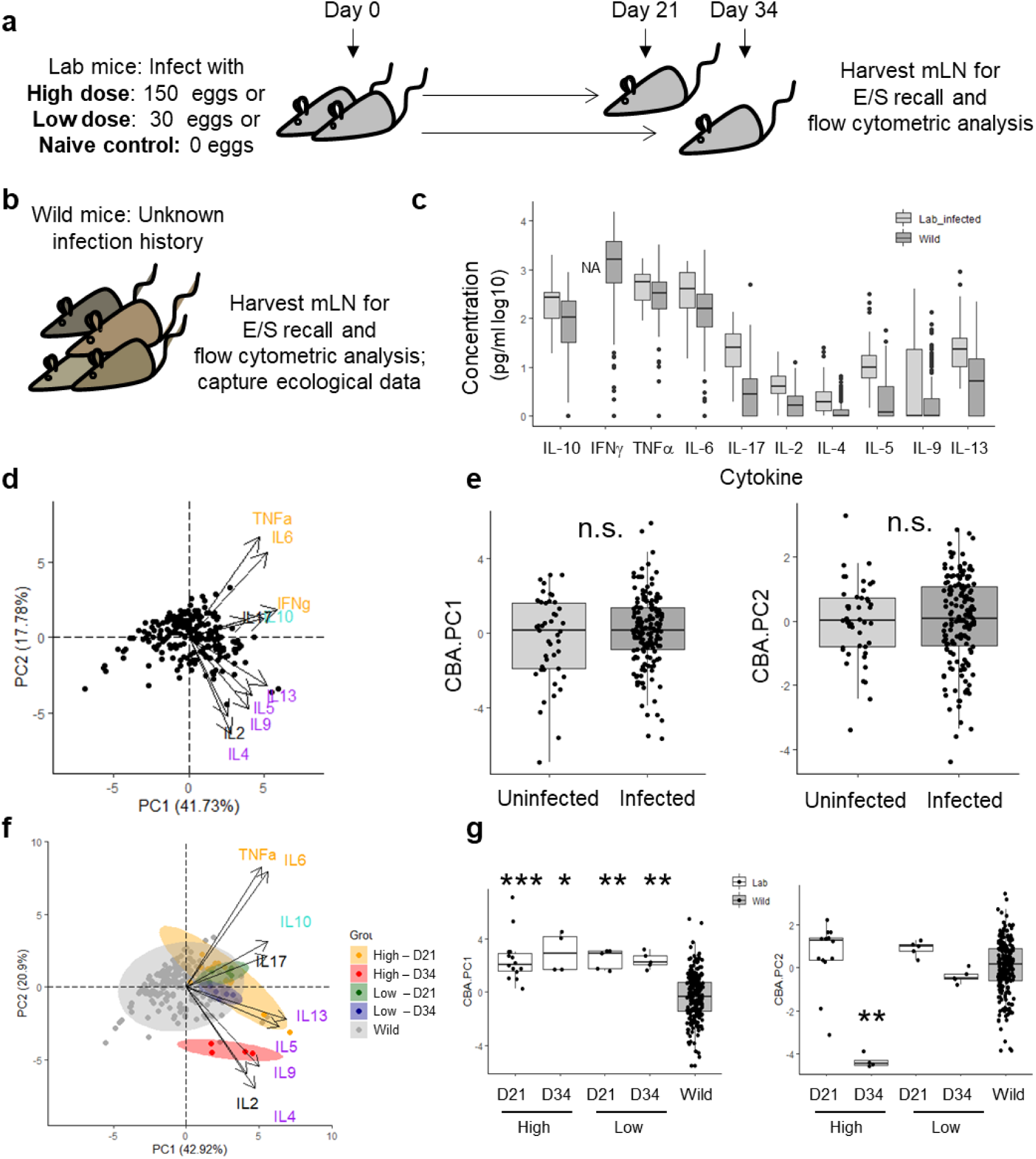
Wild mice vary in their local anti-*Trichuris* Th1/Th2 balance and secrete lower levels of cytokines than laboratory mice. **a)** Experimental design for *T. muris* infection in laboratory mice. **b)** Experimental design in wild mice. **c)** Cytokine concentrations in cell supernatants of mesenteric lymph node cells cultured for 48h in the presence of *T. muris* E/S for 48h. **d)** Principal components 1 (PC1) and 2 (PC2) of a principal component analysis of the parasite-specific cytokine response in wild mice. **e)** PC1 and PC2 of PCA analysis from (D) in uninfected versus infected wild mice. **f)** Principal components 1 (PC1) and 2 (PC2) of a principal component analysis of the parasite-specific cytokine response in wild mice and laboratory mice experimentally infected with *T. muris*. **g)** PC1 and PC2 of PCA analysis from (F) grouped by experimental group and analysed using a Kruskal-Wallis test followed by a Dunn’s test with Bonferroni adjustment. Statistical significance only presented for comparison of wild versus laboratory groups. *p≤0.5, **p≤0.01, ***p≤0.001.

### CD4^+^ T effector/memory cell phenotype is linked to antigen-specific cytokine production in wild mice but does not explain differences between wild and laboratory immune responses

Since the main source of cytokines in the recall response are CD4^+^ T cells, we next performed flow cytometric analysis to assess whether different T cell phenotypes explained the observed variation in cytokine expression in mLNs of wild animals (Suppl. Figure S5 (gating), Figure 3A). The proportion of T_EM_ within the CD4^+^ T cell pool was the only parameter positively correlated with PC1 (cytokine level) (Figure 3B), out of five parameters included in the GLM (see Suppl. Table 2 for full model output). Thus, animals with a generally heightened T cell activation status in the mLN, as determined by flow cytometry, produced more cytokines in response to an antigen-specific challenge. PC2 (Th1/Th2 balance) was significantly associated with higher proportions of T-bet and lower proportions of GATA-3 expressing effector Th1 and Th2 cells respectively, following the expected Th1/Th2 paradigm (33).

**Figure 3:**
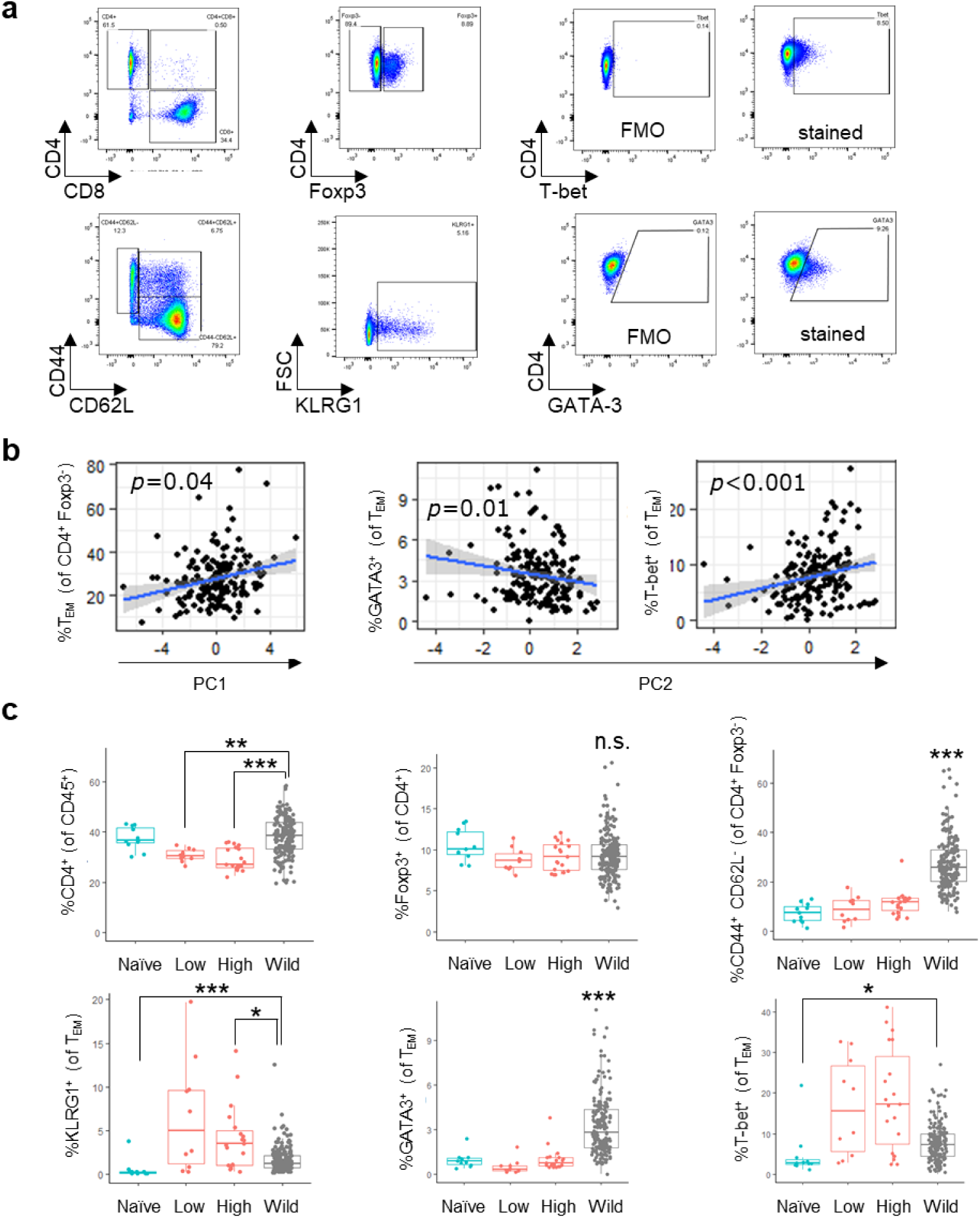
CD4^+^ effector T cell phenotype is linked to cytokine production but does not explain differences in anti-Trichuris cytokine response between laboratory and wild mice. **a)** Representative flow cytometry plots showing gating strategies for CD4^+^ T cells, Foxp3^+^ regulatory T cells, effector memory CD4^+^ T cells (T_EM_), and T-bet and GATA3 expression in T_EM,_ in mesenteric lymph nodes (mLN) of wild mice. **b)** Correlation plots of PC1 and PC2 (from Fig 2D) with selected flow cytometric mLN CD4^+^ T cell phenotypes; p values are derived from generalised linear models explaining either PC1 or PC2 with proportion of CD4+ T cells, Foxp3^+^ regulatory T cells, T_EM_, T-bet^+^ T_EM_ and GATA3^+^ T_EM_. **c)** Group-wise comparison of selected mLN CD4+ T cell phenotypes in wild (grey) and laboratory mice (blue: naive; red: infected with low or high dose of *T.muris*) using a Kruskal-Wallis test followed by a Dunn’s test with Bonferroni adjustment. Statistical significance only presented for comparison of wild versus laboratory groups. *p≤0.5, **p≤0.01, ***p≤0.001.

To check whether differential cytokine secretion could be explained by underlying differences in the cellular make-up of the mesenteric lymph nodes, we also compared the relative percentages of CD4^+^ T cell subsets in wild and laboratory mice. Lower cytokine expression levels in wild mice was not due to lower proportion of CD4^+^ or T_EM_ present in mLN, or higher Treg proportions compared to laboratory mice (Figure 3C). In fact, wild mouse mLNs harboured significantly enlarged populations of T_EM_ compared to laboratory mice. The co-inhibitory receptor KLRG1 was expressed in small proportions of T_EM_ both in laboratory infected and wild mice; thus we saw no indication of heightened T cell senescence or inhibition in the wild (34). Interestingly, we observed a Th2 propensity in wild mice based on markedly higher proportions of GATA-3 expressing T_EM_ compared to naïve or infected C57BL/6 mice, although this was not reflected by enhanced Th2 cytokine production (see Figure 2E, F). In contrast, T-bet^+^ Th1 cells were enriched in wild mice only in comparison to naive laboratory mice. In infected laboratory mice, proportions of Th1 cells were highly variable, a pattern which was associated with a temporal boost in Th1 cells at day 21 (median: 21.3; IQR: 17, 33.3) which subsided back to near naive levels by day 34 following infection (median: 3.62; IQR: 2.43, 4.44), as previously reported on the level of T helper IFN-γ expression (35, 36). Differing CD4^+^ T cell phenotypes exhibited by laboratory and wild animals were therefore insufficient to explain the differences in anti-*Trichuris* cytokine responses between laboratory and wild mouse groups.

### *Trichuris* burden and age are linked with the parasite-specific immune response in wild mice

Given the multiple host-intrinsic and host-extrinsic variables at play within our wild mouse population we asked whether certain ecological variables could explain the variation in cytokine response. A redundancy analysis (RDA), correcting for year of capture, showed that the selected explanatory variables all together explained 4.3% of the variation in cytokine expression (global RDA significance p = 0.001, see Suppl. Table 3 for full model output). Nevertheless, worm burden and maturity index were significant contributors to variation in cytokine expression (Figure 4A). Importantly, higher worm burdens were associated with strong, regulated Th1 responses (higher secretion of TNF-α, IL-6, IFN-γ, and IL-10). In contrast, increasing maturity explained the enhanced Th2 response (Figure 1D). The other variables including sex, body condition, and month of capture were not able to explain cytokine expression patterns.

**Figure 4:**
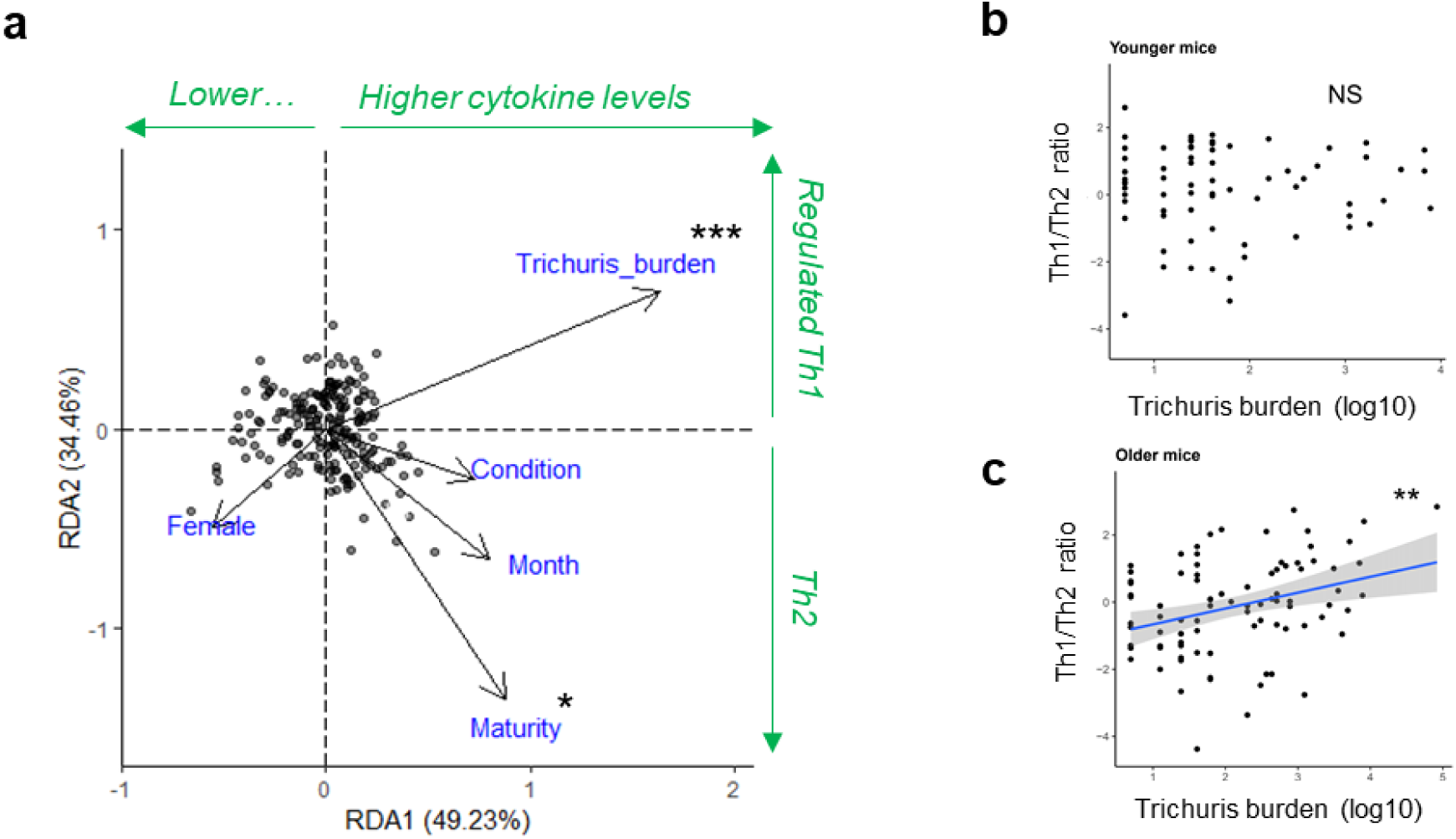
Worm burden and age contribute to variation in anti-*Trichuris* immune response. **a)** Redundancy analysis explaining variation in anti-Trichuris cytokine response in the mLN with ecological factors: Sex, maturity index as proxy for age, month of capture, scaled mass index as proxy for body condition, and *Trichuris* worm burden; with year and trap site accounted for. **b-c)** Scatterplot showing relationship between *T. muris* infection burden in infected individuals and relative strength of Th1 cytokine expression by mesenteric lymph node T cells measured by cytokine bead array, following stimulation with *T. muris* excretory-secretory molecules (using PC2 from Figure 2D). This data was separated into 2 equal cohorts of (b) lower and (c) higher maturity scores (age proxy). Explanatory terms provided to mixed model include *T. muris* burden (log counts+1), SMI, date as fixed factors, and sex as a random factor (NS=non-significant, *p≤0.5, **p≤0.01, ***p≤0.001).

The paradoxical finding that worm burdens generally increased with age (see Figure 1D), yet resistant Th2 responses also significantly increased with age warranted further investigation. Dividing the mice into equally-sized ‘younger’ and ‘older’ cohorts showed that in younger mice, the Th1/Th2 balance did not distinguish between higher and lower levels of worm burden between individuals (Figure 4B). However, strikingly, in older individuals there was a positive association of worm burden with Th1 responses (mixed model *p*=0.0028) (Figure 4B, C, see Suppl. Table 4 for full model output). Thus, on the other hand, stronger Th2 responses were associated with lower worm burdens – in the older cohort only.

### Longitudinal data highlights temporal associations between host health, infection dynamics and immunity

A powerful tool in ecological studies is the use of longitudinal data to assess the relationship between host-intrinsic and -extrinsic factors over time. Using two key metrics for host health, body condition (SMI) and gut responsiveness, we aimed to assess potential health impacts of infection longitudinally. These two metrics were negatively associated (Spearman correlation, *p*=2.27×10^−6^), and as such these analyses reflect different aspects of health in potential trade-off. We leveraged longitudinal data of body condition, faecal egg counts, and serum anti-*Trichuris* IgG1 and IgG2a levels, of a subset of 91 mice, to probe associations between changes in these values and measures of animal health (see Suppl. Figure S6 for raw data distribution). As with the worm counts, egg burden values showed an overdispersed distribution, though in individuals with both worms and eggs, there was little association between cull egg burden and worm burden (Spearman correlation, *p*=0.755, n=22). Body condition change was negatively correlated with egg burden change, in that animals with the strongest increase in body condition displayed the greatest reduction in egg burden over the course of a month, whilst animals losing body condition were more likely to display increased egg burden a month later (mixed model estimate = −0.73, *p*=0.006) (Figure 5A). Gut responsiveness showed no association with changes in egg burden (mixed model *p*=0.29), but did show a positive association with increases in anti-*T. muris* IgG1 (mixed model estimate=0.37, *p*=0.008). (Figure 5B, and for full model outputs see Suppl. Table 5).

**Figure 5:**
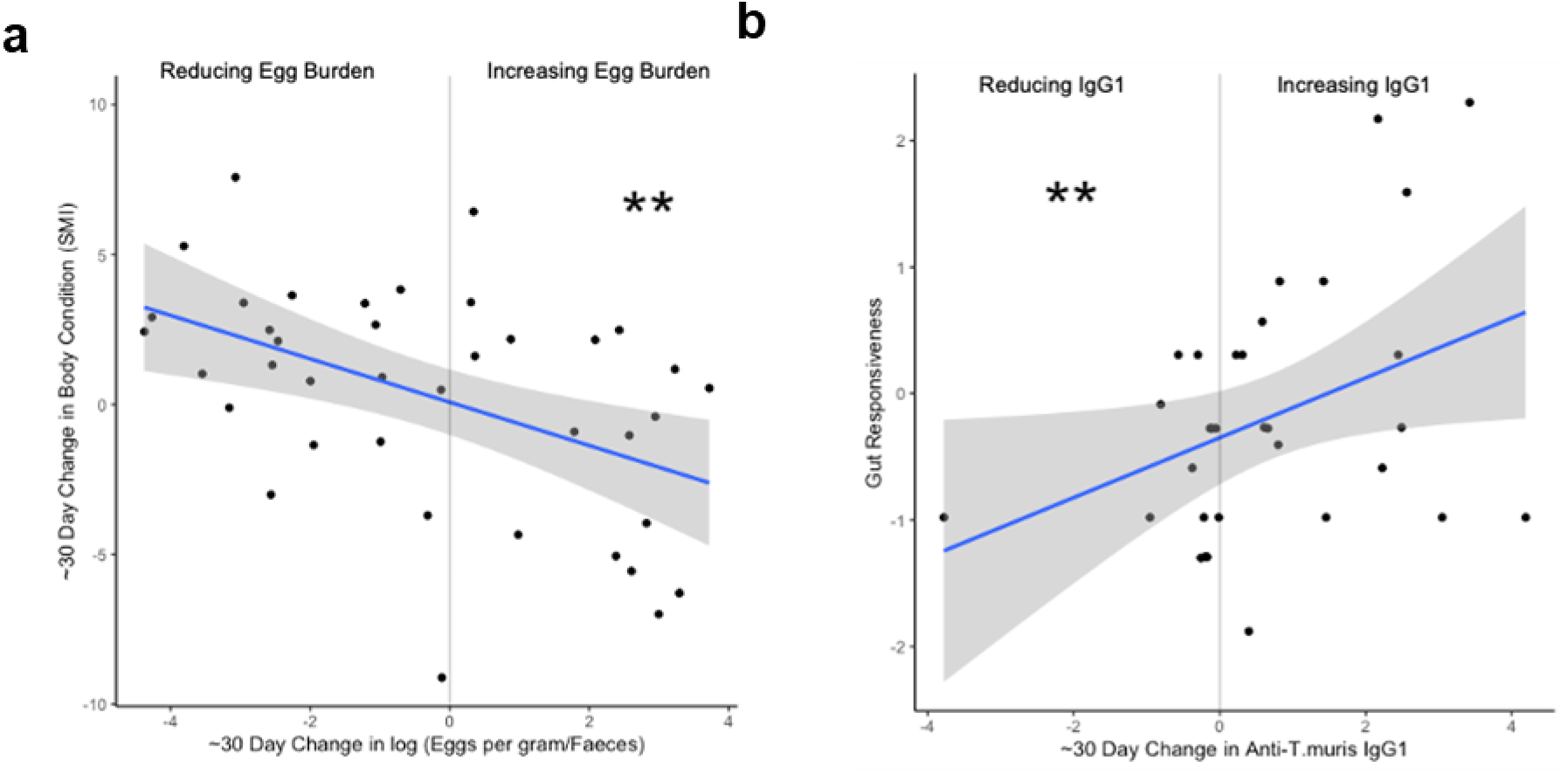
Longitudinal shifts in infection status are linked to host health. **a)** Scatter plot showing association between change in *T. muris* egg burden over 25-35 days and corresponding change in scaled mass index (SMI) as a measure of body condition (Mixed-effects model estimate=-0.73, p=0.0054). **b)** Scatter plot showing association between change in anti-*T. muris* IgG1 over 25-35 days and cull gut responsiveness score (Mixed-effects model estimate=0.37, p=0.0085). Mixed-effects models considered only individuals with changing egg burdens, and predictors included change in egg burden, change in anti-*T. muris* IgG1 and IgG2a levels, with sex and age cohort as random effects (*p≤0.5, **p≤0.01, ***p≤0.001).

## Discussion

Using a robust and extensive combination of immunological measurements and ecological data from over 200 animals, we have captured a comprehensive picture of the mammalian immune response to a natural infection with the parasitic nematode *Trichuris muris*. The prevalence of infection (>70%) in this island population of house mice is much higher than reported in many other wild rodent studies (37–39) but reflects that observed in some human populations where *T. trichiura* is endemic (17, 40). Th1/Th2-dependent immunity to *Trichuris* has been characterised in depth in laboratory mice (23, 41, 42), particularly using the C57BL/6 strain (25, 27, 35, 43). In order to better understand how well the laboratory based paradigms of resistance and susceptibility to helminth infection reflect immune responses in animals living in a multivariate environment, we compared our wild mouse data to that generated in C57BL/6 mice. Cross-sectional immunological analyses support general patterns of immunity observed from laboratory models, particularly a propensity for cytokine responses to vary along a Th1/Th2 axis, and a clear link between CD4^+^ T_EM_ cell phenotype and cytokine production. Novel insights include an age-associated dichotomy in the Th1/Th2 balance linked to difference in worm burden, as well as an overall reduced cytokine expression in response to parasite antigen compared to laboratory models.

The strength (cytokine levels) and quality (Th1/Th2 bias) were the major descriptors of variation in the *Trichuris*-specific immune response in wild mice, linked to the presence of effector T cells and their expression of T-bet/GATA3 in the mesenteric lymph nodes. Whilst the Th1/Th2 paradigm is well studied in laboratory parasite models, only few field studies have been able to confirm this immunological paradigm on a cellular level (9, 12, 14, 16, 17). Cytokine responses in the context of *Trichuris* infection in mice – albeit not parasite-specific – have only been reported in two small-scale studies in natural (14) or semi-natural settings (44). These studies point towards infected wild or re-wilded mice being more prone towards a Type 1 immune response, with heightened proportions of T-bet^+^ cells and IFNγ production being reported. Indeed, the study by Zhang et al. found no evidence of Th2 effector cells in mice infected with *Trichuris*, potentially – as the authors note themselves – due to the small size of the study cohort. In contrast, in the present study, we found evidence of a large pool of GATA3^+^ effector T cells in the gut draining mesenteric lymph nodes, at levels higher than that seen in infected laboratory mice. Nevertheless, both cytokine responses and parasite epidemiological data from this study suggest that *T. muris* infection in the Isle of May mouse population is mainly of a chronic nature and sterile immunity is rare. Older mice overall harboured more worms than younger mice, as has also been reported in a wild *Apodemus sylvaticus* population (45). In laboratory mice, ageing and immune senescence has been implicated in enhanced pro-inflammatory, Th1-dominated responses leading to higher susceptibility to infection (29). Yet, we observe an overall increase in Th2 responses with age. It is noteworthy, however, that the majority of the mouse population on the Isle of May lives for less than one year due to population crashes each winter (46), and our study population therefore focuses on immune changes seen between juveniles and early adulthood, rather than senescence. Of the ecological variables tested to explain variation in cytokine response in wild mice, only age and worm burden came out as significant, despite causative links of sex (28) and high-fat diet (27) with the immune response to this parasite in laboratory settings. These results showcase that experimental effects seen in laboratory conditions can be dampened or superseded by the abundance of other variation at play in the wild. Strikingly, in younger wild animals, Th1/Th2 balance was varied but did not associate with worm burden. However, in older wild animals, a link between Th2 responses and lower worm burden did emerge, as expected from laboratory studies. Studies in humans infected with soil transmitted helminths, including the human whipworm *T. trichiura*, have also described significant negative associations between chronic worm abundance and Th2 immune responses in an age-dependent manner (16, 17). Adopting the approach of dividing the study population into two age classes, both studies revealed that only in the older cohort, stronger Th2 immune responses were associated with reduced susceptibility, in keeping with our Isle of May study. Thus, it appears that in younger individuals in the wild – whether mouse or human – immune polarisation per se is not enough to control worm burden; only with increasing age, potentially through repeated infection exposure or changes in other host-associated factors, the Th1/Th2 balance becomes a determinant of susceptibility or resistance. Interestingly, in voles, it has been reported that older, but not younger animals display a tolerant phenotype of increasing worm burden concomitant with increasing body condition, linked with expression of GATA3 on a whole-blood RNA level(47). This age-associated dichotomy has not been captured in laboratory mice thus far, where young animals are routinely used and display a clear link between Th1/Th2 balance and disease susceptibility. We would argue that in fact it may be hard to capture this dichotomy in comparatively small laboratory mouse studies due to the homogeneity of age, environmental variables, and exposure regimes. The Isle of May mouse population thus provides a tractable real world model for dissecting the underlying immune mechanisms at play in determining resistance and susceptibility to infection.

Overall, we observed a reduced cytokine response in wild compared to laboratory mice in response to parasite-specific stimulation of mLN cells, despite an enlarged CD4^+^ T_EM_ pool compared to laboratory mice, typical for wild or microbially-exposed laboratory mice (13, 48–50). Significantly lower cytokine responses in wild mice have been observed in response to innate immune stimulation, but not consistently in response to stimulation of the adaptive immune response (13, 49). Likewise, exposure of laboratory mice to wild mouse microbiome (48, 51, 52) or environmental antigens (53) have pointed towards a heightening of the adaptive immune response compared to clean laboratory mice. Searching for the underpinning mechanism, we did not find evidence of enhanced T cell senescence, an immunological mechanism by which effector T cell responses to the infection could be curtailed (54).

Whilst laboratory models cannot capture the complexity of the real world and work with immature immune systems(48, 55), our data still support the relevance of laboratory mouse systems in serving as models for animal populations living in uncontrolled settings. Thus, wild mice did not cluster entirely separately from all laboratory mice; overlap was observed with laboratory animals currently harbouring infection. The Th1/Th2 bias of the immune response in wild mice was comparable to that of single bolus low-dose infected laboratory mice, despite low level infections in the wild mice likely the consequence of the ingestion of low numbers of eggs over time (56). Laboratory infection models have attempted to replicate infection scenarios analogous to those assumed to occur in natural conditions for soil-transmitted helminths. *Trichuris* trickle infection experiments revealed that slow accrual of small numbers of worms over time enabled an eventual threshold of worm burden to be reached, at which point the mouse switched from a more Th1-dominated to a Th2 phenotype accompanied by worm loss, in a highly genotype-dependent manner (25, 57). The precise control offered over the time of infection and its subsequent progression is of course also a strength of the laboratory model. For example, we were unable to distinguish between parasite-specific cellular immune responses mounted by currently infected hosts versus currently uninfected (but seropositive) hosts. The variable infection history and concomitant ecological variation appeared to supersede the difference in immune response expected from a currently infected and a currently uninfected individual.

In addition to our cross sectional data we also present a longitudinal infection dataset to highlight dynamic shifts occurring between an animal’s health and infection status. Whilst immunomodulatory parasite products are being explored for their potential therapeutic effect in autoimmune and allergic disease (58), it is clear that chronic STH infections can cause a significant health burden in affected individuals (15). Time-series egg-count datasets have been used to elucidate fitness trade-offs associated with immunity to helminth infection and predict survival through crash years in wild mammals (59, 60). While longitudinal analyses can ideally make the directionality of these immune dynamics clearer, the results from analyses of changes between two time points showed little association between immunity and parasite fecundity, as indicated by egg shedding, over time. The fact that egg counts are not a strong indicator of worm burdens limits the usefulness of these analysis if egg counts are used as a proxy for worm burden, and should be kept in mind for other field studies using egg burdens as an indicator of infection levels. We suggest that egg burden could be taken as a ‘worm fecundity’ measure, and whilst it is known that egg shedding can vary throughout the day, repeated measures still provide a valuable insight into parasite fitness. The increase in body condition concomitant with a decrease in faecal egg burdens over time indicates that infection may confer some level of fitness cost in the mice, with measurable health benefits of reducing worm fecundity (60–62). The positive association observed between increasing levels of anti-*T.muris* IgG1 and gut responsiveness at cull suggests that driving of IgG1 levels, likely an indicator of stronger Th2 dominated immune responses (63), may come at the cost of local tissue damage. Th2 immune responses can induce both tissue regenerative and pathological processes (64); especially in the wild, it is likely that trade-offs are at play between immunity and associated energetic costs and tissue damage (65). Further longitudinal work, with multiple time points per animal would help to elucidate the directionality of relationships between immunity, body condition and infection.

To conclude, laboratory mice and wild mice differ in multiple ways each of which can influence their immune “phenotype”. For example, laboratory mice do not have to contend with natural pressures such as temperature shifts, food scarcity, reproduction, or co-infections (66), and the immune system of laboratory animals is immature in phenotype and function compared to ‘dirtier’ animals (48, 50, 51, 53, 55). Diversifying experimental parameters such as sex and age, microbial exposure of laboratory mice, outdoor housing of laboratory mice, or the use of fully wild mouse populations, are therefore key additional approaches to investigating the immune system in order to develop a clear understanding of its complexity and function (2). Studies in a wild sheep population have demonstrated that endoparasite worm burdens predict winter survival (60) and in human populations there is clear evidence of reduced quality of life in individuals infected with *Trichuris trichiura* (15). What constitutes an appropriate immune response to parasites to minimise adverse effects on health – either due to exacerbated immune investment or due to costs of parasitism – and how these immune responses are shaped, are two of the big outstanding questions to be addressed in free-living or re-wilded populations.

## Methods

### Mice

Wild house mice were live-trapped between August and December in 2018 and 2019 on the Isle of May (56°11′11·6″N, 2°33′24·1″W) on three independent trapping grids of 96 Longworth traps, each placed 8–10 metres apart in a 6 × 16 grid and containing Sizzle Nest (Datesand, catalogue number CS1A09) and sunflower seeds, as described in Mair et al. 2021 (67). Male C57BL/6 mice were bought from Envigo and maintained at a temperature of 20–22°C in a 12-h light–12-h dark lighting schedule, in sterile, individually ventilated cages in same-sex groups of 2–5, with food and water ad lib. Laboratory mice were 8–13 weeks old when used for this study. All animals used for this study were euthanized by a rising concentration of CO_2_.

### *Trichuris* DNA extraction and PCR sequencing

Genomic DNA from 3 female worms belonging to the Isle of May and Edinburgh *T. muris* isolates (6 in total) were extracted via a DNeasy Blood and Tissue kit (Qiagen) as per the manufacturer’s instructions. Quality of DNA extractions were determined via a 0.8% agarose gel stained with SYBR™ Safe (Thermofisher). The ITS1-5.8S-ITS2 rDNA regions of each worm was then amplified via PCR using the nematode specific NC5: GTAGGTGAACCTGCGGAAGGATCATT and NC2: TTAGTTTCTTTTCCTCCGCT primer set. Each PCR included 0.5 µl of each primer (0.5 µM), 12.5 µl of OneTaq Quick-Load 2X Master Mix with Standard Buffer (New England Biolabs), 2 µl of diluted (1:10) genomic DNA, and 10 µl of MilliQ water. PCR cycling parameters consisted of initial denaturation at 94°C for 5 mins, followed by 35 cycles of 94°C for 30 seconds, 54°C for 30 seconds, and 72°C for 60 seconds, accompanied by a final elongation step of 72°C for 10 minutes. Obtained PCR products where then isolated on a 1% agarose gel stained with SYBR™ Safe (Thermofisher). DNA bands pertaining to the amplified PCR products were cut out of the gel under UV light and extracted from the agarose via a Qiaquick gel extraction kit (Qiagen) according to the manufacturer’s protocols. Concentrations of the extracted PCR products were quantified via a Qubit 4 flurometer. Subsequently, obtained PCR products were diluted to 40 ng/µl in 10 µl of MilliQ water and underwent Sanger sequencing performed by the Genomic Technologies Core Facility at the University of Manchester.

### *Trichuris* phylogenetic analysis

All phylogenetic analysis was performed in MEGA11. ITS1-5.8S-ITS2 rDNA sequences of nematode species were aligned, and outlying regions were removed to make sequences uniform in length (977 bp) except for the distantly related species *Ascaris lumbricoides* and *Trichuris mastomysi* which were used to root the phylogenetic trees. The most appropriate phylogenetic model for each alignment of sequences was determined via the “*Find Best DNA/Protein Models (ML)”* function provided in MEGA11. For the sequence alignment underpinning Suppl. Figure S1, which contained ITS1-5.8S-ITS2 rDNA sequences pertaining to the *T. muris* Isle of May isolate alongside numerous other *Trichuris* species the maximum Likelihood method in tandem with the Tamura-Nei model and a discrete Gamma distribution to model evolutionary rate differences among sites (T92+G) was recommended. In contrast, only the maximum Likelihood method in conjunction with the Tamura-Nei model (T92) was suggested for the alignment associated with Suppl. Figure S2 that consisted of many different *T. muris* haplotypes dispersed across Europe, published in Callejón et al. 2010 (31) and expanded by Wasimuddin *et al*. (68).

### Infection of laboratory mice with *Trichuris muris*

Mice were infected with *T*. *muris* eggs via oral gavage in a final volume of 200 μl of deionized water. For low-dose *T*. *muris* infection, 30 embryonated eggs were given to each mouse, and for high-dose infection, 100 infective embryonated eggs were given. Parasite maintenance, assessing of egg infectivity and counting of eggs were performed as described previously (69).

### Tissue preparation and cell isolation

Mesenteric lymph nodes were dissected and manually dissociated through 70 μm filters. Cells were counted using haemocytometers and 0.4% nigrosin (Sigma-Aldrich) dilutions for the exclusion of dead cells on the Isle of May, and using a CASY cell counter (Scharfe System) at the University of Manchester. Cells were re-suspended at 1×10^7^ cells/ml in RPMI 1640 buffer supplemented with 10% foetal calf serum (FCS, Sigma-Aldrich) for further analysis. The gastrointestinal (GI) tract was manually dissected, and a 0.3-0.4 cm snip of proximal colon fixed overnight in 10% neutral buffered formalin (Fisher), followed by long-term storage in 70% ethanol. Another 0.3-0.4 cm snip was fixed overnight in methacarn (64% methanol, 27% chloroform, 9% acetic acid), then transferred into 100% methanol for 30 minutes followed by 100% ethanol until processing. The residual gastrointestinal tissue was stored in 80% ethanol for later parasitological analysis.

### Mesenteric lymph node re-stimulation and cytokine bead array

Mesenteric lymph node cells were cultured in flat-bottomed 96 well plates at 5×10^6^ cells/ml for 48h at 37°C 5% CO_2_ in RPMI 1640 supplemented with 10% FCS, 1% L-Glutamine, and 1% penicillin/streptomycin (Invitrogen), with 50μg/ml of excretory-secretory product (prepared as detailed in (69). Supernatants were harvested and stored at −20°C until cytokine determination. Cytokines were analysed using the Cytometric Bead Array (CBA) Mouse/Rat soluble protein flex set system (BD Bioscience), which was used according to the manufacturer’s instructions. Bead acquisition was performed on a MACSQuant (Miltenyi Biotech). For cytokine concentration analysis, FCAP Array v3.0.1 software (BD Cytometric Bead Array) was used. 10 cytokines were quantified in this assay: IFNγ, TNFα, IL-2, IL-4, IL-5, IL-6, IL-9, IL-10, IL-13 & IL-17.

### Flow cytometry

Single-cell suspensions from mesenteric lymph nodes were washed thoroughly with PBS and stained with Fixable Viability Dye eFluor 455UV and anti-CD16/CD32 (both from Thermo Fisher) in PBS for 10 min prior to addition of the relevant surface marker fluorochrome-conjugated antibodies in FACS buffer (PBS, 2% heat-inactivated FCS) supplemented with Super Bright staining buffer (Thermo Fisher). Following 30 min of incubation, cells were washed with FACS buffer and fixed Fix-Perm buffer of the eBioscience™ Foxp3 / Transcription Factor Staining Buffer Set (Thermo Fisher) overnight. Cell suspensions were kept at 4°C and in the dark throughout incubation steps. The following morning, samples were washed in Perm buffer of the same Buffer Set and stained with relevant intracellular marker fluorochrome-conjugated antibodies in Perm buffer for 30min at room temperature. After a final wash with Perm buffer, samples were re-suspended in FACS buffer and kept at 4°C in the dark until acquisition within 6 days of staining. The following antibodies were used: Ki-67-ef450 (clone SOLA15), Foxp3-ef506 (clone FJK-16S), CD62L-SB600 (clone MEL-14), CD44-SB645 (clone IM7), CD19-SB702 (clone EBIO1D3), CD4-SB780 (clone RM4-5), ICOS-FITC (clone C398.7), CD8-PerCP-ef710 (clone 53-6.7), ST2-PE (clone RMST2-33), KLRG1-PE-ef610 (clone 2F1), GATA-3-PE-Cy7 (clone TWAJ), T-bet-ef660 (clone 4B10), CD45-Af700 (clone 104), CD11b-APC-ef780 (clone M1/70). Samples were acquired on an LSRFortessa running FACSDiva 8 software (Becton Dickinson, Wokingham, UK). Data were analysed using FlowJo software (TreeStar; version 10.4.2). For gating strategy see Suppl. Figure S5.

### Serum antibody detection

Blood samples were taken from the tail vein for longitudinal sampling and/or after euthanasia via cardiac puncture. Serum was retrieved by centrifugation of clotted blood samples and stored at −80°C prior to processing. Parasite-specific serum levels of IgG1 and IgG2c antibody were measured via ELISA. 96 well plates were coated with 5μg/ml *T. muris* overnight excretory-secretory product in PBS overnight, plates were washed, and non-specific binding blocked with 3% BSA (Sigma-Aldrich) in PBS. Plates were washed and incubated with serum dilutions (for IgG1 1:10,000; for IgG2 1:5,000) in triplicates. Each well also had a standard curve in duplicates using a single serially-diluted and aliquoted serum sample from a pool of T. muris infected laboratory mice across the entire study, allowing for inter-plate variability to be controlled for. Parasite specific antibody was measured using biotinylated IgG1 (BD Biosciences) or IgG2c (BD Biosciences) antibodies which were detected with SA-POD (Roche). After a final wash, plates were developed with TMB substrate kit (BD Biosciences, Oxford, UK) according to the manufacturer’s instructions. The reaction was stopped using 0.18 M H_2_SO_4_ after 10 min incubation in the dark. The plates were read by a Versa max microplate reader (Molecular Devices) through SoftMax Pro 6.4.2 software at 450 nm, with reference of 570 nm subtracted. For each standard concentration on each plate, the mean OD value was calculated and a 4-parameter logistic model was fit across the serial dilution to form a standard curve for each plate. These standard curves were then used to estimate concentrations from OD values for the mouse blood serum wells. From sample triplicates, the greatest outlier was removed and the average of the remaining two values calculated. These averaged concentration values were then used for any downstream analysis.

### Parasitology

Prior to dissection of the ethanol-fixed gastrointestinal (GI) tract, the liquid in the tube was examined in a scored petri dish under dissecting microscope, to count any worms which had moved out of the GI tract during storage and transportation. The GI tract was then separated into stomach, small intestine, caecum and colon. Each section was opened in a scored petri dish using dissection tools, and gently washed out with water to dislodge any material inside. The washed-out contents, and the tissue lining, were examined under dissection microscope, and any worms counted.

Faecal samples were collected during live trapping and at cull, and transferred directly to 1.6ml tubes containing sodium acetate-acetic acid-formalin (SAF) solution. Egg counts were made using a modified version of the McMaster flotation method. Faecal samples were homogenised by manual disruption with a bent metal seeker, and then centrifuged at 400G for 10 minutes. The proportion by volume of the resulting pellet to the liquid SAF component was recorded. For each sample, a glass slide was divided into 10 equal sections. The pellet was again manually disrupted and homogenised, and in turn, 10μl of homogenised sample was pipetted across each slide section and checked under a microscope at ×10 magnification. The number of *T. muris* eggs per section were counted and averaged. The average egg count per section and SAF-to-pellet volume ratio were used to calculate number of eggs per ml. All parasitological surveys were conducted at the University of Nottingham.

### Gut responsiveness score

A ‘gut responsiveness score’ was derived based on variation in colonic crypt length, extent of inflammatory infiltrate, depth of inflammatory infiltrate, and goblet cell numbers per crypt in wild mice. We chose an alternative phrase to ‘gut pathology score’, standardly used in laboratory colitis models or human IBD samples, as we cannot ascertain whether a higher score is a pathological response in wild animals.

Neutral-buffered-formalin fixed tissues were dehydrated through a graded series of ethanol, cleared in xylol and infiltrated with paraffin in a dehydration automat (Leica ASP300 S) using a standard protocol. Specimens were embedded in paraffin (Histocentre2, Shandon), sectioned on a microtome (5μm sections) and allowed to dry for a minimum of 4 hours at 40°C. Prior to standard H&E staining, slides were deparaffinised with citroclear (TCS biosciences) and rehydrated through alcohol (100% to 70%) to PBS or water. H&E stained sections were analysed using CaseViewer; crypt length was measured, and extent (graded 0-4) and depth (mucosal, submucosal, transmural) of inflammatory infiltrate assessed.

In order to measure goblet cells per crypt, methacarn-fixed tissues were cleared in xylol and infiltrated with paraffin in a dehydration automat (Leica ASP300 S) using a standard protocol prior to paraffin-embedding and tissue sectioning as describe above. Mucins in goblet cells were stained with 1% alcian blue (Sigma-Aldrich) in 3% acetic acid (Sigma-Aldrich, pH 2.5) for 5 mins. Sections were washed and treated with 1% periodic acid, 5 minutes (Sigma-Aldrich). Following washing, sections were treated with Schiff’s reagent (Vicker’s Laboratories) for 10 minutes and counterstained in Mayer’s haematoxylin (Sigma-Aldrich). Slides were then dehydrated and mounted in DPX mounting medium (Thermo Fisher). For enumeration of goblet cell staining, the average number of cells from 20 crypts was taken from three different sections per mouse. Images were acquired on a 3D-Histech Pannoramic-250 microscope slide-scanner using a *20x/ 0*.*30 Plan Achromat* objective (Zeiss).

Four measures of gut responsiveness (crypt length, extent of inflammatory infiltrate, depth of inflammatory infiltrate, and goblet cell numbers per crypt) were used in a PCA to examine their covariance. PC1, explaining 43.4% variation, described a positive association between all 4 variables (Suppl. Figure S4). The PC1 eigenvalues for each individual were extracted for use as a multiparameter ‘gut responsiveness score’ in downstream analyses.

### Assessment of ecological variables and calculation of proxies

Sex of the animal was assessed visually at cull. Scaled Mass Index (70) was calculated from body length and mass as a measure of body condition relative to the whole population. While age is typically estimated from dry eye lens mass in wild rodent populations, building on the formulae set out by Rowe *et al*. 1985 (71), we found this method to not be fully accurate on its own, as ages were calculated which were known to be impossible i.e. mice were known to be older than the calculated ages based on recapture data. In order to develop a useful and continuous proxy measure of age, a principal component analysis (PCA) was performed on cull samples, incorporating body mass, body length, tail length and dry paired eye lens mass, in order to provide a ‘maturity index’. PC1 explained 69.6% of variation in these samples, and PC1 scores were taken as maturity index values. In order to have an equivalent index for non-cull samples, another PCA was carried out across all samples, excluding eye lens mass. PC1 explained 69.3% of variation in this analysis. These maturity PCA scores were strongly correlated with the analogous scores from cull (linear model p<2.2^−16^, R^2^=0.95), and so the longitudinal maturity index was taken as an appropriate measure of biological age when required for longitudinal analysis. Further PCA details and loadings are provided in Supporting Materials.

## Statistical analyses

General: All data were analysed using R version 4.1.2 (2021-11-01) (72). Relevant statistical tests are described in results sections and figure legends, and for generalised linear models the full model outputs are included in supplementary tables as indicated.

Redundancy analysis: To explain how patterns of variation in cytokine profiles are explained by infection with *T. muris*, against a variable ecological background, redundancy analysis were performed on cytokine concentration data from the cytokine bead array, using the R package ‘**vegan**’ (Oksanen et al, 2020). Cytokine concentration values were used as response variables, and *T. muris* infection burdens (log (count+1) values) SMI, trapping month, age (as maturity index estimate), and sex were included as explanatory variables, with year and trap site accounted for (n = 199). The redundancy model was tested for overall significance, as well as each axis of variation, and each explanatory variable.

Longitudinal analysis: For longitudinal analysis, a subset of the dataset was used for which two time points of faecal egg counts were available, and which had associated anti-*T. muris* antibody concentration data for both time points (n=87). For all analyses involving egg counts, log (count+1) values were used. Values of change in SMI and anti-*T.muris* IgG1 and IgG2a were calculated (henceforth ΔSMI, ΔIgG1, and ΔIgG2a, respectively). Two mixed models were performed with change in body condition, and gut responsiveness at cull included as response variables. As we specifically aimed to model change in egg burden, we excluded any individuals with no eggs present in faecal samples at both time-points (remaining n = 40). Included as independent variables were change in log(eggs per gram + 1), ΔIgG1 and ΔIgG2a,, with sex and age cohort (where maturity index <0 = “Young” and maturity index >0 = “Mature”) as random factors.

## Abbreviations

CBA: Cytokine bead array
E/S: Excretory-secretory product
FACS: Fluorescence-activated cell sorting
FCS: Foetal calf serum
FEC: Feacal egg count
GI tract: Gastrointestinal tract
GLM: Generalised linear model
GLMM: Generalised linear mixed model
mLN: Mesenteric lymph nodes
PBS: Phosphate-buffered saline
PCA: Principal component analysis
PC1 / 2: First two principal components of the principal component analysis
RDA: Redundancy analysis
RDA1 / 2: First two dimensions of the ordination space from the redundancy analysis
SMI: Scaled mass index
STH: Soil-transmitted helminths
T_EM_: Effector/memory CD4^+^ T cells
Treg: CD4^+^ Foxp3^+^ regulatory T cells

## Acknowledgements

We would like to thank Nature Scotland for permission to carry out work on the Isle of May; David Steel (Nature Scotland), Bex Outram (Nature Scotland) and Mark Newell (Centre for Ecology and Hydrology) for support with fieldwork; and our many fieldwork volunteers. We thank Prof. John Brookfield (University of Nottingham) and Prof. Andrew MacColl (University of Nottingham) for their feedback on the manuscript drafts. We thank the University of Manchester Flow Cytometry Core Facility and the Genomic Technologies Core Facility for their technical support.

## Author contributions

JEB and KJE conceived the overarching project and should be considered joint last authors. KJE and IM designed laboratory experiments. AW and JEB designed the fieldwork. AW, AL and IM organized the fieldwork. IM, AW, AL, JEB, KJE, AB, AM and JF collected field data. IM, AB, JT and LL performed laboratory experiments. JF, IM and OD analysed the data. IM, JF, KJE, and JEB interpreted the data. IM and JF led the writing of the paper, with significant feedback and contribution from SS. IM and JF contributed equally to this publication and should be considered joint first authors. All authors contributed critically to the drafts and gave final approval for publication.

## Ethical approval

The work on wild mice was approved by the University of Nottingham Animal Welfare and Ethical Review Body and complies with the UK’s Animals (Scientific Procedures) Act of 1986. The work on laboratory mice was approved by the University of Manchester Local Animal Welfare and Ethical Review Body and complies with the UK’s Animals (Scientific Procedures) Act of 1986.

## References

1. L. K. Beura, et al., Normalizing the environment recapitulates adult human immune traits in laboratory mice. Nature 532, 512–516 (2016).

2. I. Mair, T. N. McNeilly, Y. Corripio-Miyar, R. Forman, K. J. Else, Embracing nature’s complexity: Immunoparasitology in the wild. Seminars in Immunology 53, 101525 (2021).

3. S. A. Budischak, et al., Feeding immunity: Physiological and behavioral responses to infection and resource limitation. Frontiers in Immunology 8, 8 (2018).

4. F. Yeung, et al., Altered Immunity of Laboratory Mice in the Natural Environment Is Associated with Fungal Colonization. Cell Host and Microbe 27, 809–822.e6 (2020).

5. R. K. Boughton, G. Joop, S. A. O. Armitage, Outdoor immunology: methodological considerations for ecologists. Functional Ecology 25, 81–100 (2011).

6. A. B. Pedersen, S. A. Babayan, Wild immunology. Molecular Ecology 20, 872–880 (2011).

7. J. E. Bradley, J. A. Jackson, Measuring immune system variation to help understand host-pathogen community dynamics. Parasitology 135, 807–823 (2008).

8. S. Young, et al., Relationships between immune gene expression and circulating cytokine levels in wild house mice. Ecology and Evolution 10, 13860–13871 (2020).

9. A. J. Vo Ezenwa, Opposite effects of anthelmintic treatment on microbial infection at individual versus population scales. Science 347, 175–177 (2015).

10. E. Arriero, et al., From the animal house to the field: Are there consistent individual differences in immunological profile in wild populations of field voles (Microtus agrestis)? PLOS ONE 12, e0183450 (2017).

11. J. A. Jackson, et al., An Immunological Marker of Tolerance to Infection in Wild Rodents. PLOS Biology 12, e1001901 (2014).

12. Y. Corripio-Miyar, et al., Functionally distinct T-helper cell phenotypes predict resistance to different types of parasites in a wild mammal. Scientific Reports 2022 12:1 12, 1–12 (2022).

13. S. Abolins, et al., The comparative immunology of wild and laboratory mice, Mus musculus domesticus. Nat. Commun. 8, 14811 (2017).

14. H. Zhang, L. Bednář, E. Heitlinger, S. Hartmann, S. Rausch, Whip- and pinworm infections elicit contrasting effector and distinct regulatory responses in wild house mice. International Journal for Parasitology 52, 519–524 (2022).

15. K. J. Else, et al., Whipworm and roundworm infections. Nature Reviews Disease Primers 6 (2020).

16. J. D. Turner, et al., Th2 Cytokines Are Associated with Reduced Worm Burdens in a Human Intestinal Helminth Infection. Journal of Infectious Diseases 188, 1768–1775 (2003).

17. J. A. Jackson, et al., Cytokine response profiles predict species-specific infection patterns in human GI nematodes. International Journal for Parasitology 34, 1237–1244 (2004).

18. D. A. P. Bundy, E. S. Cooper, D. E. Thompson, J. M. Didier, I. Simmons, Epidemiology and population dynamics of Ascaris lumbricoides and Trichuris trichiura infection in the same community. Transactions of the Royal Society of Tropical Medicine and Hygiene 81, 987–993 (1987).

19. N. A. Croll, E. Ghadirian, Wormy persons: Contributions to the nature and patterns of overdispersion with Ascaris lumbricoides, Ancylosotma duodenale, Necator americanus and Trichuris trichiura. Tropical and Geographical Medicine 33, 241–248 (1981).

20. J. M. Behnke, J. W. Lewis, S. N. M. Zain, F. S. Gilbert, Helminth infections in Apodemus sylvaticus in southern England: Interactive effects of host age, sex and year on the prevalence and abundance of infections. Journal of Helminthology 73, 31–44 (1999).

21. T. I. A. Roach, D. Wakelin, K. J. Else, D. A. P. Bundy, Antigenic cross-reactivity between the human whipworm, Trichuris trichiura, and the mouse trichuroids Trichuris muris and Trichinella spiralis. Parasite Immunology 10, 279–291 (1988).

22. B. J. Foth, et al., Whipworm genome and dual-species transcriptome analyses provide molecular insights into an intimate host-parasite interaction. Nature Genetics 2014 46:7 46, 693–700 (2014).

23. K. J. Else, F. D. Finkelman, C. R. Maliszewski, R. K. Grencis, Cytokine-mediated regulation of chronic intestinal helminth infection. The Journal of experimental medicine 179, 347–351 (1994).

24. D. Sorobetea, M. Svensson-Frej, R. Grencis, Immunity to gastrointestinal nematode infections. Mucosal Immunology 11, 304–315 (2018).

25. A. J. Bancroft, K. J. Else, N. E. Humphreys, R. K. Grencis, The effect of challenge and trickle Trichuris muris infections on the polarisation of the immune response. International Journal for Parasitology 31, 1627–1637 (2001).

26. E. Michael, D. A. P. Bundy, The effect of the protein content of CBA/Ca mouse diet on the population dynamics of Trichuris muris (Nematoda) in primary infection. Parasitology 103, 403–411 (1991).

27. E. Funjika, et al., High-fat diet-induced resistance to helminth infection via alternative induction of type 2 immunity. Mucosal Immunology 16, 27–38 (2023).

28. M. R. Hepworth, M. J. Hardman, R. K. Grencis, The role of sex hormones in the development of Th2 immunity in a gender-biased model of Trichuris muris infection. European Journal of Immunology 40, 406–416 (2010).

29. N. E. Humphreys, R. K. Grencis, Effects of ageing on the immunoregulation of parasitic infection. Infection and Immunity 70, 5148–5157 (2002).

30. J. E. Keeling, Experimental trichuriasis. I. Antagonism between Trichuris muris and Aspiculuris tetraptera in the albino mouse. The Journal of parasitology 47, 641–646 (1961).

31. R. Callejón, et al., Molecular evolution of Trichuris muris isolated from different Muridae hosts in Europe. Parasitology Research 107, 631–641 (2010).

32. S. Goertz, et al., Geographical location influences the composition of the gut microbiota in wild house mice (Mus musculus domesticus) at a fine spatial scale. PLoS ONE 14 (2019).

33. M. Ruterbusch, K. B. Pruner, L. Shehata, M. Pepper, In Vivo CD4+ T Cell Differentiation and Function: Revisiting the Th1/Th2 Paradigm. 10.1146/annurev-immunol-103019-085803 38, 705–725 (2020).

34. S. M. Henson, A. N. Akbar, KLRG1—more than a marker for T cell senescence. AGE 2009 31:4 31, 285–291 (2009).

35. K. S. Hayes, A. J. Bancroft, R. K. Grencis, The role of TNF-α in Trichuris muris infection I: influence of TNF-α receptor usage, gender and IL-13. Parasite Immunology 29, 575–582 (2007).

36. K. J. Else, R. K. Grencis, Cellular immune responses to the murine nematode parasite Trichuris muris. I. Differential cytokine production during acute or chronic infection. Immunology 72, 508 (1991).

37. M. Ito, T. Itagaki, Survey on Wild Rodents for Endoparasites in Iwate Prefecture, Japan. Journal of Veterinary Medical Science 65, 1151–1153 (2003).

38. M. Fasihi Harandi, S. M. Madjdzadeh, M. Ahmadinejad, Helminth parasites of small mammals in Kerman province, southeastern Iran. Journal of Parasitic Diseases 40, 106–109 (2016).

39. P. D. Carrera-Játiva, et al., Gastrointestinal parasites in wild rodents in Chiloé Island-Chile. Revista Brasileira de Parasitologia Veterinária 32, e017022 (2023).

40. L. Hakami, et al., Epidemiology of soil transmitted helminth and Strongyloides stercoralis infections in remote rural villages of Ranomafana National Park, Madagascar. Pathogens and Global Health 113, 94 (2019).

41. K. J. Else, G. M. Entwistle, R. K. Grencis, Correlations between worm burden and markers of Th1 and Th2 cell subset induction in an inbred strain of mouse infected with Trichuris muris. Parasite Immunology 15, 595–600 (1993).

42. A. J. Bancroft, K. J. Else, R. K. Grencis, Low-level infection withTrichuris muris significantly affects the polarization of the CD4 response. European Journal of Immunology 24, 3113–3118 (1994).

43. N. E. Humphreys, R. K. Grencis, Effects of ageing on the immunoregulation of parasitic infection. Infection and Immunity 70, 5148–5157 (2002).

44. J. M. Leung, et al., Rapid environmental effects on gut nematode susceptibility in rewilded mice. PLoS Biol 16, e2004108 (2018).

45. J. M. Behnke, F. S. Gilbert, M. A. Abu-Madi, J. W. Lewis, Do the helminth parasites of wood mice interact? Journal of Animal Ecology 74, 982–993 (2005).

46. G. S. Triggs, The population ecology of house mice (Mus domesticus) on the Isle of May, Scotland. Journal of Zoology 225, 449–468 (1991).

47. J. A. Jackson, et al., An Immunological Marker of Tolerance to Infection in Wild Rodents. PLOS Biology 12, e1001901 (2014).

48. L. K. Beura, et al., Normalizing the environment recapitulates adult human immune traits in laboratory mice. Nature 532, 512–516 (2016).

49. S. R. Abolins, M. J. O. Pocock, J. C. R. Hafalla, E. M. Riley, M. E. Viney, Measures of immune function of wild mice, Mus musculus. Molecular Ecology 20, 881–892 (2011).

50. A. S. Japp, et al., Wild immunology assessed by multidimensional mass cytometry. Cytometry Part A 91, 85–95 (2017).

51. S. P. Rosshart, et al., Wild Mouse Gut Microbiota Promotes Host Fitness and Improves Disease Resistance. Cell 171, 1015–1028.e13 (2017).

52. S. P. Rosshart, et al., Laboratory mice born to wild mice have natural microbiota and model human immune responses. Science 365 (2019).

53. H. Arnesen, et al., Naturalizing laboratory mice by housing in a farmyard-type habitat confers protection against colorectal carcinogenesis. Gut Microbes 13 (2021).

54. L. Haynes, A. C. Maue, Effects of aging on T cell function. Current Opinion in Immunology 21, 414–417 (2009).

55. S. P. Rosshart, et al., Laboratory mice born to wild mice have natural microbiota and model human immune responses. Science 365 (2019).

56. J. M. Behnke, D. Wakelin, The survival of Trichuris rnuris in wild populations of its natural hosts. Parasitology 67, 157–164 (1973).

57. M. Glover, S. A. P. Colombo, D. J. Thornton, R. K. Grencis, Trickle infection and immunity to Trichuris muris. PLoS Pathogens 15, 1–27 (2019).

58. R. M. Maizels, H. H. Smits, H. J. McSorley, Modulation of Host Immunity by Helminths: The Expanding Repertoire of Parasite Effector Molecules. Immunity 49, 801–818 (2018).

59. A. L. Graham, et al., Fitness correlates of heritable variation in antibody responsiveness in a wild mammal. Science 330, 662–665 (2010).

60. J. A. Leivesley, et al., Survival costs of reproduction are mediated by parasite infection in wild Soay sheep. Ecology Letters 22, 1203–1213 (2019).

61. S. A. Budischak, D. O’Neal, A. E. Jolles, V. O. Ezenwa, Differential host responses to parasitism shape divergent fitness costs of infection. Functional Ecology 32, 324–333 (2018).

62. K. J. Vandegrift, T. R. Raffel, P. J. Hudson, PARASITES PREVENT SUMMER BREEDING IN WHITE-FOOTED MICE, PEROMYSCUS LEUCOPUS. Ecology 89, 2251–2258 (2008).

63. K. A. Mowen, L. H. Glimcher, Signaling pathways in Th2 development. Immunological Reviews 202, 203–222 (2004).

64. R. L. Gieseck, M. S. Wilson, T. A. Wynn, Type 2 immunity in tissue repair and fibrosis. Nature Reviews Immunology 2017 18:1 18, 62–76 (2017).

65. M. Zuk, A. M. Stoehr, Immune defense and host life history. American Naturalist 160 (2002).

66. A. L. Graham, Naturalizing mouse models for immunology. Nat Immunol 22, 111–117 (2021).

67. I. Mair, et al., A lesson from the wild: The natural state of eosinophils is Ly6Ghi. Immunology 164, 766–776 (2021).

68. Wasimuddin, et al., Testing parasite “intimacy”: The whipworm Trichuris muris in the European house mouse hybrid zone. Ecology and Evolution 6, 2688–2701 (2016).

69. R. Forman, et al., Trichuris muris infection drives cell-intrinsic IL4R alpha independent colonic RELMα+ macrophages. PLOS Pathogens 17, e1009768 (2021).

70. J. Peig, A. J. Green, New perspectives for estimating body condition from mass/length data: the scaled mass index as an alternative method. Oikos 118, 1883–1891 (2009).

71. F. P. Rowe, A. Bradfield, R. J. Quy, T. Swinney, Relationship Between Eye Lens Weight and Age in the Wild House Mouse (Mus musculus). The Journal of Applied Ecology 22, 55 (1985).

72. R Core Team, R: A language and environment for statistical computing. (2021).

