## Supplementary Figures and Tables for "The adaptive immune response to *Trichuris* in wild versus laboratory mice: An established model system in context"

| Term | Variance | p |
|---|---|---|
| SMI | -4.83E-03 | 0.882 |
| Maturity | 0.194 | **2.55E-05*** ** |
| Sex | -0.152 | 0.32 |
| Date | -5.25E-06 | 0.999 |

**Supplementary Table 1. Summary statistics of *T. muris* worm burden GLM**

Each row corresponds to an explanatory variable provided to a generalised linear model, using log(*T. muris* worm burden+1) as the response (*p≤0.5, **p≤0.01, ***p≤0.001 , n=256).

**a**

| Term | Estimate | p |
|---|---|---|
| (Intercept) | -0.3651798 | 0.7408 |
| $CD4^+$ T cells (of $CD45^+$) | -0.0262565 | 0.2223 |
| $CD44^+$ $CD62L^-$ $T_{EM}$ (of $CD4^+$ $Foxp3^-$) | 0.035137 | **0.0402*** |
| $GATA3^+$ (of $T_{EM}$) | 0.0006797 | 0.9931 |
| T-$bet^+$ (of $T_{EM}$) | -0.0339388 | 0.2931 |
| $Foxp3^+$ Treg (of $CD4^+$) | 0.0541236 | 0.4164 |
| $KLRG1^+$ (of $T_{EM}$) | 0.0905113 | 0.3824 |

**b**

| Term | Estimate | p |
|---|---|---|
| (Intercept) | 0.941735 | 0.14427 |
| $CD4^+$ T cells (of $CD45^+$) | -0.024491 | 0.05149 |
| $CD44^+$ $CD62L^-$ $T_{EM}$ (of $CD4^+$ $Foxp3^-$) | -0.005337 | 0.59021 |
| $GATA3^+$ (of $T_{EM}$) | -0.112694 | **0.01438*** |
| T-$bet^+$ (of $T_{EM}$) | 0.061975 | **0.00117**** |
| $Foxp3^+$ Treg (of $CD4^+$) | 0.028375 | 0.46437 |
| $KLRG1^+$ (of $T_{EM}$) | 0.055673 | 0.35629 |

**Supplementary Table 2. Summary statistics of flow cytometry GLM**

Each row corresponds to an explanatory variable provided to a generalised linear model, using **a)** mesenteric lymph node cytokine concentration PC1 (representing overall total concentration/strength of cytokine response) and **b)** PC2 (relative dominance of Th1 over Th2 cytokine concentration) as response variables. (*p≤0.5, **p≤0.01, ***p≤0.001 , n=162).

**a**

| | Overall Model | RDA1 | RDA2 | RDA3 | RDA4 | RDA5 |
|---|---|---|---|---|---|---|
| **Variance Explained** | 0.066 | 0.325 | 0.227 | 0.073 | 0.026 | 0.009 |
| **F** | 2.869 | 7.062 | 4.944 | 1.594 | 0.559 | 0.187 |
| **p** | **0.001*** | **0.015 *** | **0.034*** | 0.609 | 0.96 | 0.99 |
| | | **Loadings** | | | | |
| **IFN-γ** | | 0.310 | 0.091 | -0.454 | -0.238 | -0.255 |
| **IL-10** | | 0.663 | 0.127 | -0.0436 | 0.387 | -0.294 |
| **IL-13** | | 0.319 | -0.385 | -0.180 | -0.303 | 0.593 |
| **IL-17** | | -0.167 | -0.124 | -0.753 | 0.424 | 0.179 |
| **IL-2** | | -0.193 | -0.310 | -0.335 | -0.314 | -0.525 |
| **IL-4** | | 0.109 | -0.364 | 0.195 | 0.2610 | -0.028 |
| **IL-5** | | 0.376 | -0.436 | 0.143 | -0.230 | -0.237 |
| **IL-6** | | 0.250 | 0.277 | -0.054 | 0.258 | 0.026 |
| **IL-9** | | 0.144 | -0.328 | 0.039 | 0.186 | 0.264 |
| **TNF-α** | | 0.244 | 0.457 | -0.128 | -0.448 | 0.249 |

**b**

| Term | Variance | Significance |
|---|---|---|
| **SMI** | 0.081 | 0.102 |
| **Trichuris burden** | 0.024 | **0.003**** |
| **Sex** | 0.079 | 0.125 |
| **Maturity** | 0.146 | **0.017*** |
| **Month** | 0.099 | 0.067 |
| **Residual** | 8.789 | |

**Supplementary Table 3. Summary of cytokine redundancy analysis. a)** Top - Summary statistics of each redundancy axis, Bottom – loadings of each axis, showing the relative contribution of concentrations of different cytokines to each axis. **b)** Summary statistics of explanatory variables provided for redundancy analysis. *p≤0.5, **p≤0.01, ***p≤0.001. SMI; Scaled mass index.

| | Juvenile Cohort | | Mature Cohort | |
|---|---|---|---|---|
| | Estimate | p | Estimate | p |
| **(Intercept)** | 0.271 | 0.881 | 3.471 | 0.199 |
| **Trichuris burden** | 0.0481 | 0.656 | 0.359 | **0.00282**** |
| **Date** | -0.00764 | 0.0568 | -0.0106 | 0.0717 |
| **SMI** | 0.106 | 0.0394 | -0.0487 | 0.429 |

**Supplementary Table 4. Summary of age-separated cytokine-Trichuris burden mixed models.** Summary statistics for two mixed-effect models incorporating MLN cytokine PC2 (representing relative Th1 versus Th2 dominance) as the response variable, in juvenile (**left**: maturity index < 0) and mature (**right**: maturity index > 0) mice. Log(worm burden + 1), body condition (SMI) and date were included as fixed factors, and sex was included as a random factor. *p≤0.5, **p≤0.01, ***p≤0.001 , n=33. SMI; Scaled mass index.

| Response Variable: | Change in body condition (SMI) | | Gut responsiveness | |
|---|---|---|---|---|
| | **Estimate** | **p values** | **Estimate** | **p values** |
| **(Intercept)** | -0.6971 | 0.28501 | -0.44303 | 0.37511 |
| **Change in faecal egg burden** | -0.7303 | **0.00554**** | -0.0824 | 0.28661 |
| **Change in anti-*T. muris* IgG1** | 0.866 | 0.05753 | 0.3653 | **0.00845**** |
| **Change in anti-*T. muris* IgG2a** | 0.2169 | 0.62624 | -0.18901 | 0.22877 |

**Supplementary Table 5. Summary of longitudinal mixed models.** Summary statistics for two mixed-effect models incorporating longitudinal data, with the response variables as ~30 day change in body condition (left) and gut responsiveness score (right). Age cohort (where maturity index > 0 equates to the 'mature' cohort, and < 0 the 'young' cohort) and sex were included as random factors. *p≤0.5, **p≤0.01, ***p≤0.001 , n=33. SMI; Scaled mass index.

### Supplementary Figure 1

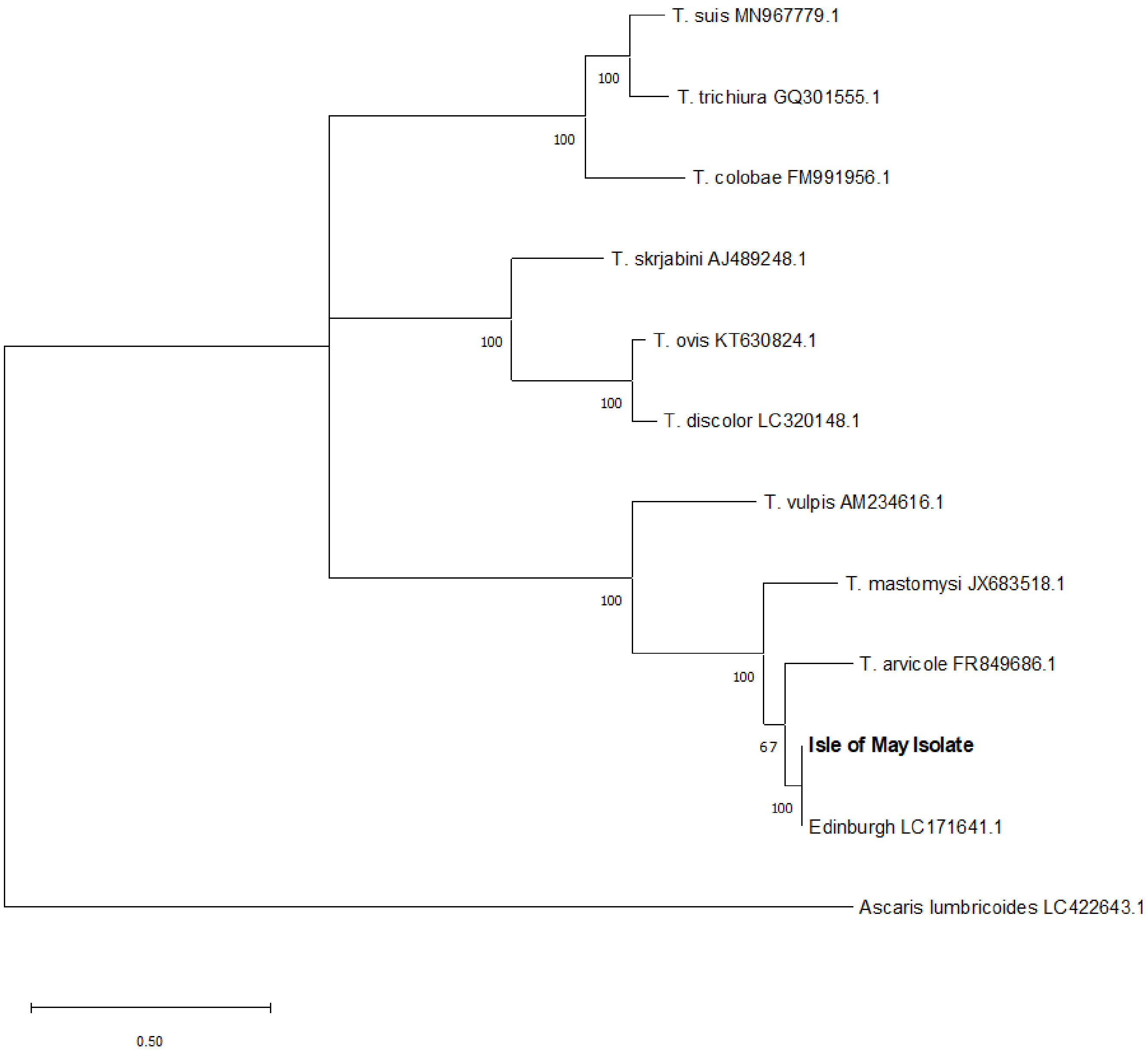


**Supplementary Figure 1. Evolutionary history of whipworms isolated from house mice on the Isle of May.** The phylogenetic tree generated by the MEGA11 software is based on the maximum Likelihood method in tandem with the Tamura-Nei model and a discrete Gamma distribution to model evolutionary rate differences among sites (T92+G) as selected by MEGA11 as the best fit model. The numbers below the branches represent nodes with >50% bootstrap support from trees generated from 1,000 bootstrap replicates and the maximum likelihood tree topology is scaled to the expected number of nucleotide substitutions per site and is defined by a scale bar (bottom left). The distantly related parasitic roundworm *Ascaris Lumbricoides* is used to root the tree. Sequences for known *Trichuris spp.* are labelled according to their species name followed by their GenBank accession number (e.g. AM234616.1). The sequence of the Edinburgh *T. muris* isolate is labelled by location and GenBank accession number (e.g. Edinburgh LC171641.1). The sequence for the unknown *Trichuris* species sampled from house mice on the isle of May is outlined in bold.

### Supplementary Figure 2

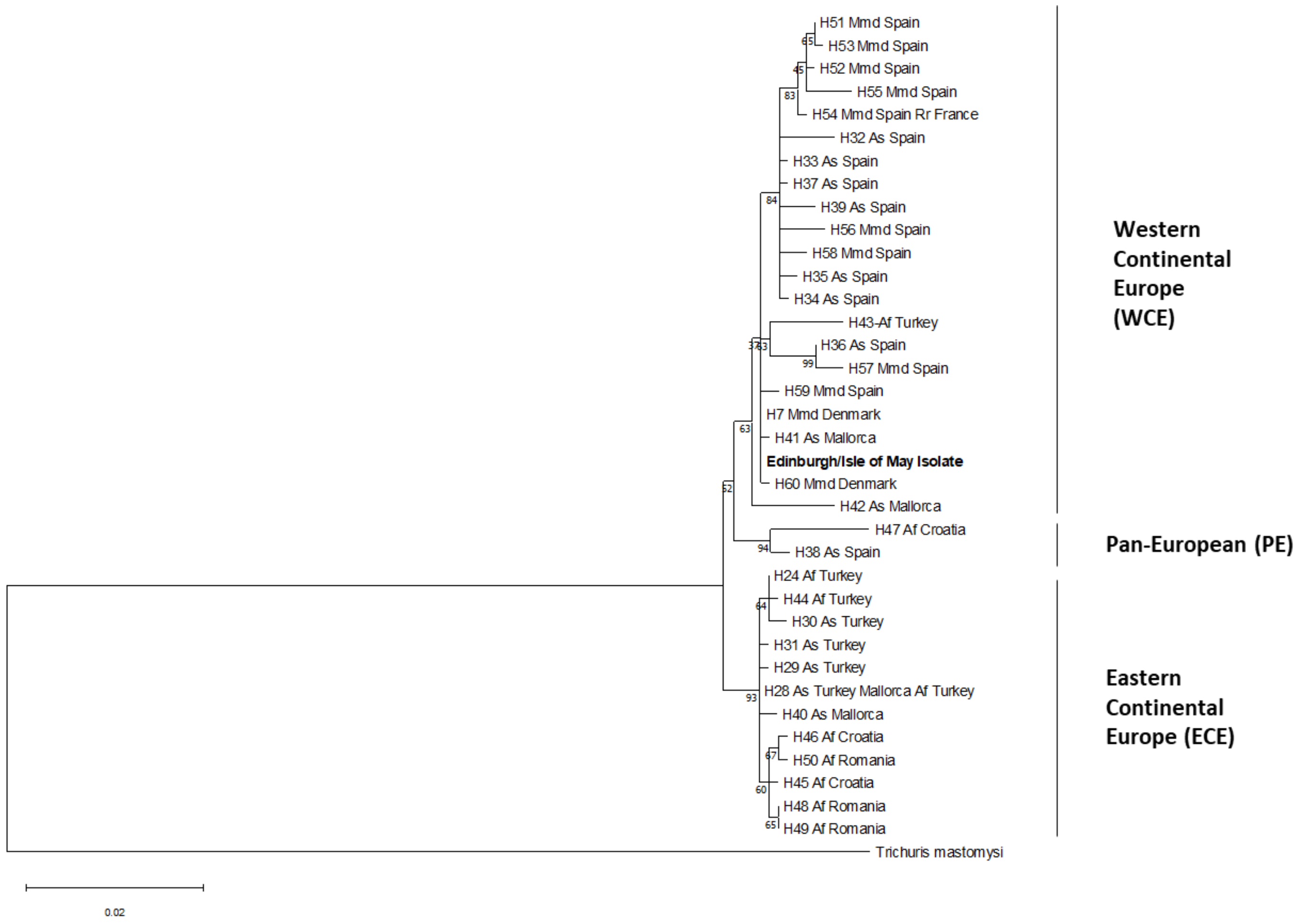


**Supplementary Figure 2. Phylogenetic analysis of the relationship between *T. muris* parasites found in house mice on the Isle of May and those sampled from house mice, rats, and wood mice throughout Continental Europe and island of Mallorca via comparison of ITS1, 5.8s, ITS2 regions in the ribosomal DNA of *T. muris* (Callejon et al. 2010).** The phylogenetic tree was generated using the MEGA11 software. The evolutionary history between the parasites was inferred using the Maximum Likelihood method in conjunction with the Tamura-Nei model (T92) as selected by MEGA11 as the best fit model. The numbers below the branches represent nodes with >50% bootstrap support from trees generated from 1,000 bootstrap replicates. *Trichuris mastomysi* was used to root the tree. The maximum likelihood tree topology is scaled to the expected number of nucleotide substitutions per site and is defined by a scale bar (bottom left). The two Continental European clusters and Pan-European cluster identified by Callejon et al. and Wasmuddin et al., respectfully, are outlined to the right of the tree. The sequence from the Isle of May/Edinburgh strain of the parasite is highlighted in bold. Sequences derived from Callejon et al. are labelled according to haplotype (e.g. H51), host species (As: Apodemus sylvaticus, Af: Apodemus flavicollis, Mmd: Mus musculus domesticus, Rr: Rattus rattus), and location (e.g. Turkey).

### Supplementary Figure 3

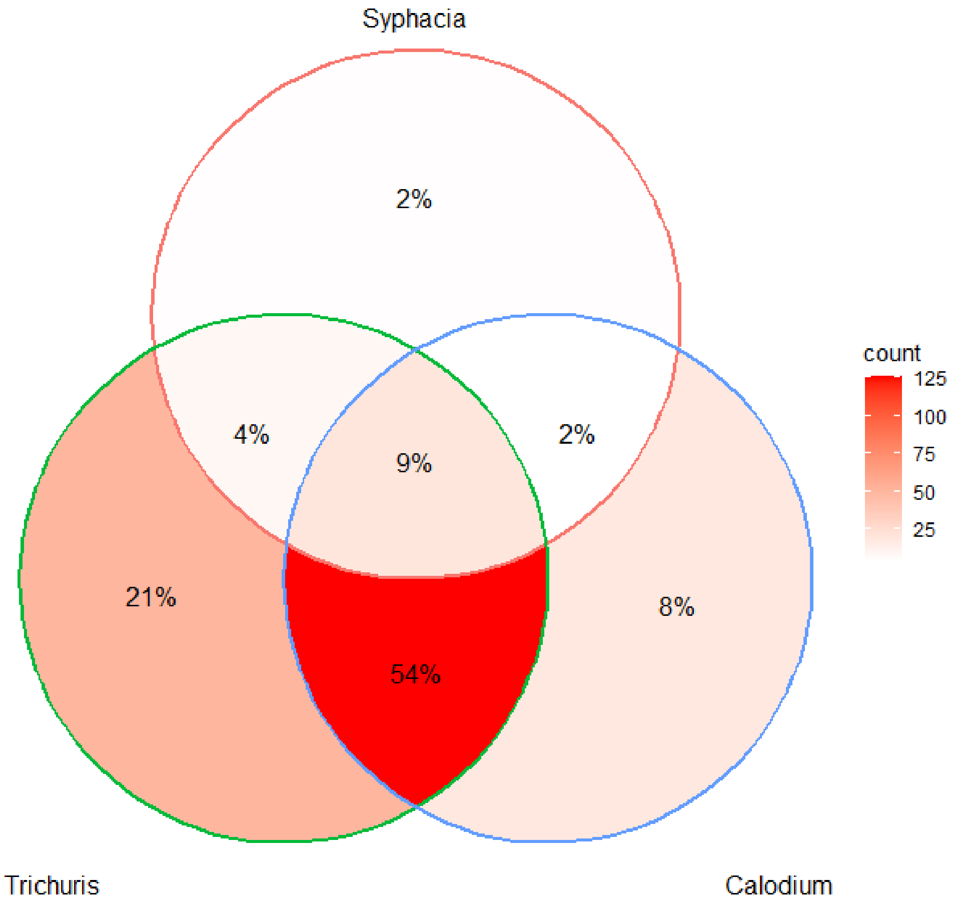


**Supplementary Figure 3. Co-infection rates for the three nematode parasite infections present in Isle of May Mice**. These include whipworm (*Trichuris muris*), pinworm (*Syphacia obvelata*) and the capillariasis-causing hepatic nematode *Calodium hepaticum*. Prevalence was confirmed through gastrointestinal dissection surveys of mature worms for *T. muris* & *S. obvelata*, and characteristic lesions and discolouration of liver tissue for *C. hepaticum*. Percentages indicate the proportion of the host population displaying a given combination of infections, with larger proportions of the population indicated by darker red colouration of the venn diagram.

### Supplementary Figure 4

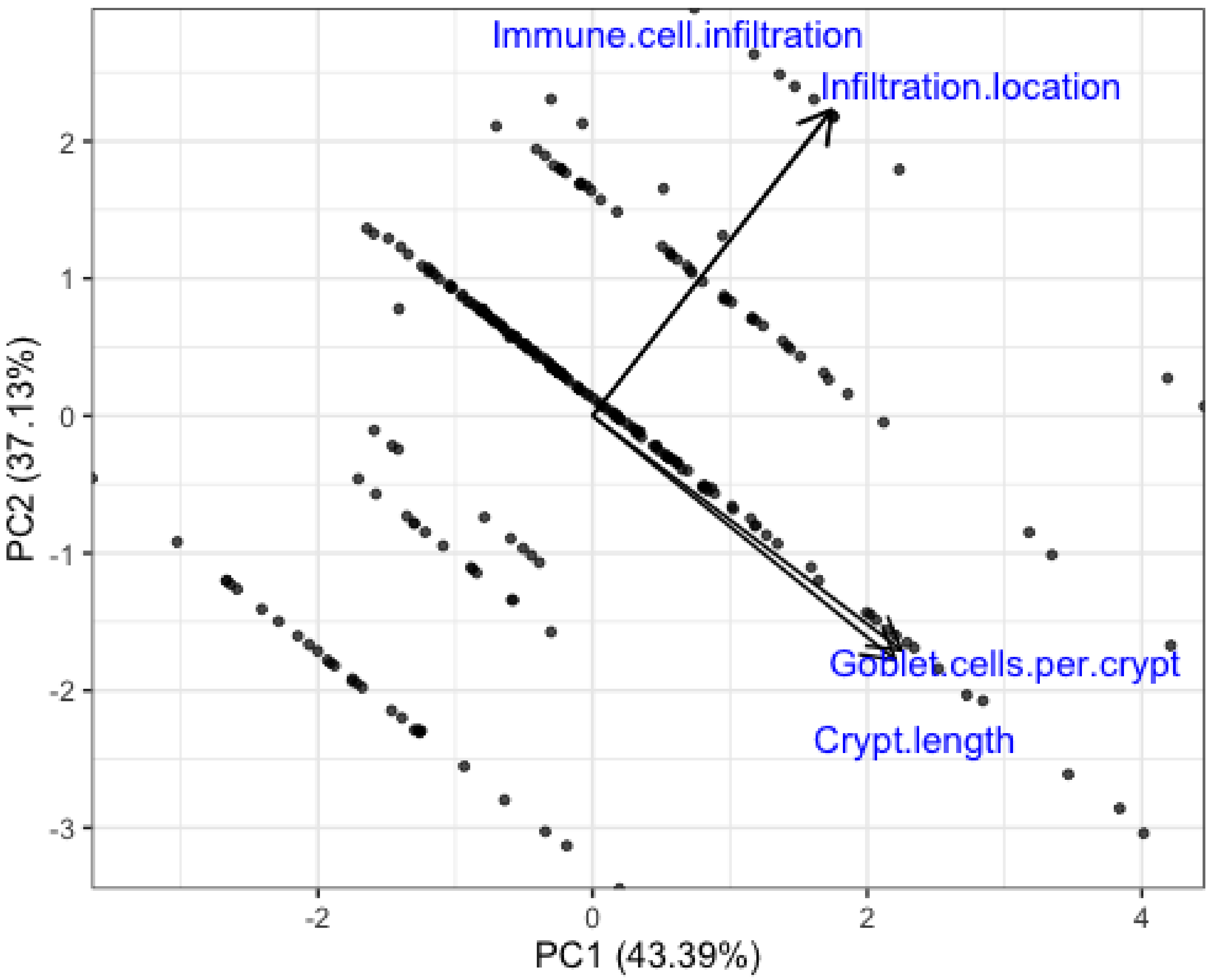


**Supplementary Figure 4. Composite score for gut responsiveness.** Ordination plot showing principal component analysis (PCA) of gut responsiveness measures. PC1 describes an increasing level of gut responsiveness across all 4 included measures, and as such PC1 scores were extracted for use as a multiparameter ‘gut responsiveness score’ in further analyses. Methods of assessment of responsiveness components are included in methods and materials.

### Supplementary Figure 5

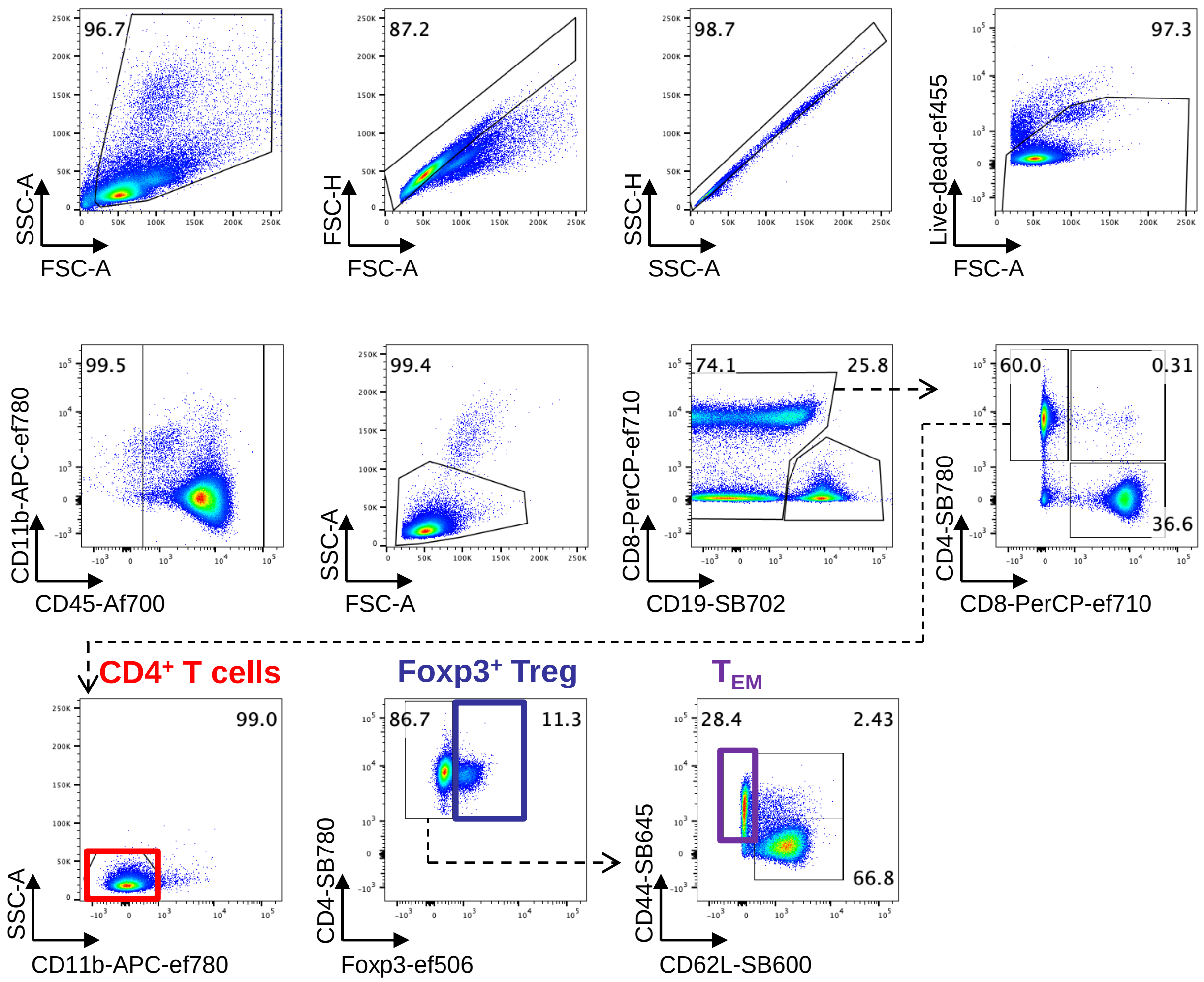


**Supplementary Figure 5. Flow cytometric gating strategy for CD4⁺ T cell subsets in wild mice.** Mesenteric lymph nodes were collected from wild house mice from the Isle of May between November 2018 and December 2019, and single cell suspensions stained for flow cytometric analysis. Representative flow cytometry plots showing gating strategy for CD4⁺ T cells (red box), Foxp3⁺ regulatory T cells (Treg, orange box), effector memory CD4⁺ T cells ($T_{EM}$) (purple box).

### Supplementary Figure 6

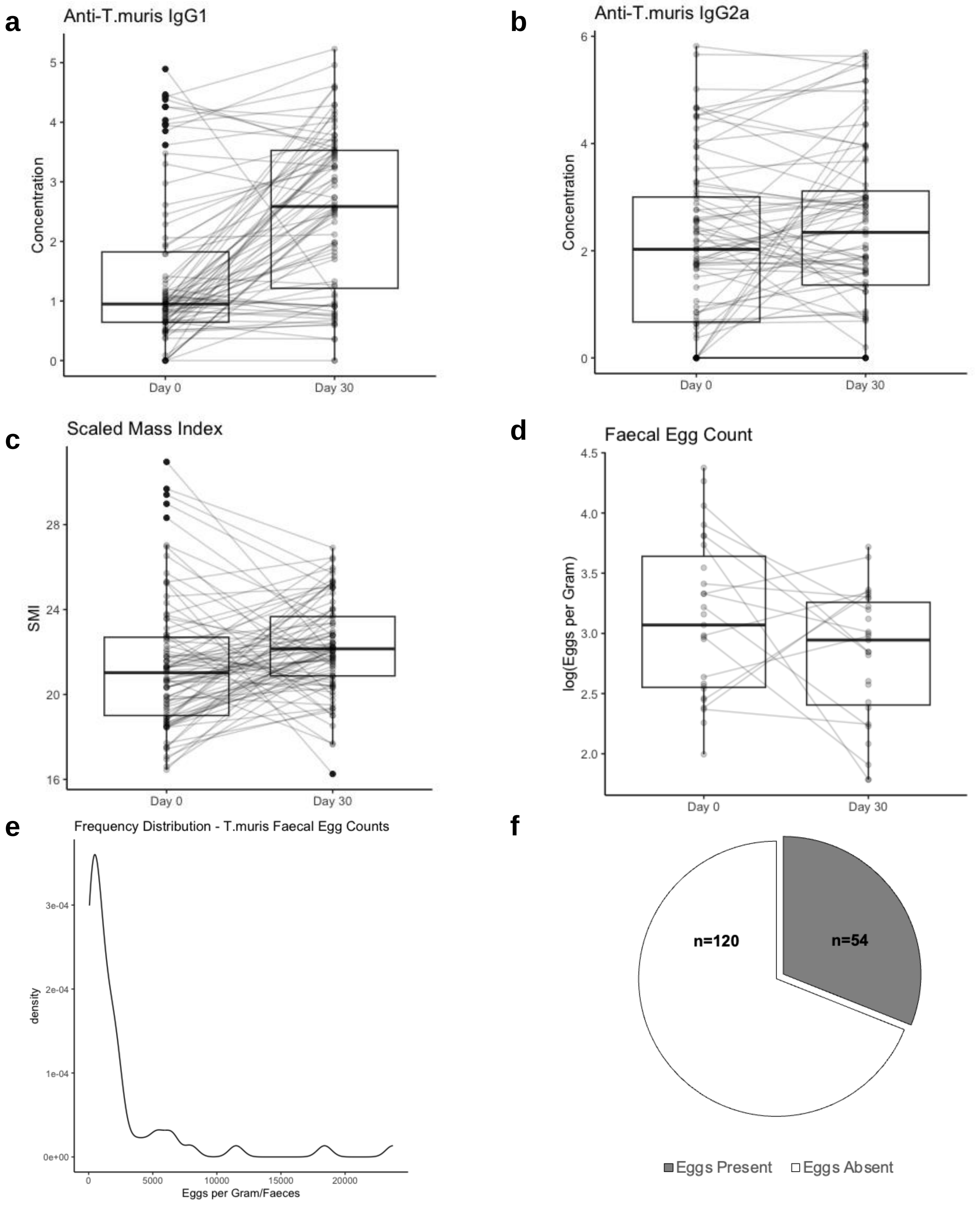


**Supplementary Figure 6. Longitudinal data distribution summary.** Paired boxplots show changes between point of initial capture to 30 days later (± 5 days) within lines connecting individual mice, across four longitudinal measures: **a)** Serum anti-*T. muris* IgG1 concentration (Box-Cox normalised), **b)** serum anti-*T. muris* IgG2a concentration (Box-Cox normalised), **c)** scaled mass index, a measure of body condition & **d)** *T. muris* eggs per gram of faeces (log10 transformed, excluding samples with zero eggs). **e)** Distribution of faecal *T. muris* egg counts (excluding samples with zero eggs). **f)** Proportion of faecal samples where *T. muris* eggs are present versus absent.
